# Molecular Interactions and Forces that Make Proteins Stable: A Quantitative Inventory from Atomistic Molecular Dynamics Simulations

**DOI:** 10.1101/2023.01.23.525230

**Authors:** Juan José Galano-Frutos, Javier Sancho

## Abstract

Protein design requires a deep control of protein folding energetics, which can be determined experimentally on a case-by-case basis but is not understood in sufficient detail. Calorimetry, protein engineering and biophysical modeling have outlined the fundamentals of protein stability, but these approaches face difficulties in elucidating the specific contributions of the intervening molecules and elementary interactions to the folding energy balance. Recently, we showed that, using Molecular Dynamics (MD) simulations of native proteins and their unfolded ensembles, one can calculate, within experimental error, the enthalpy and heat capacity changes of the folding reaction. Analyzing MD simulations of four model proteins (CI2, barnase, SNase and apoflavodoxin) whose folding enthalpy and heat capacity changes have been successfully calculated, we dissect here the energetic contributions to protein stability made by the different molecular players (polypeptide and solvent molecules) and elementary interactions (electrostatic, van der Waals and bonded) involved. Although the proteins analyzed differ in length (65-168 amino acid residues), isoelectric point (4.0-8.99) and overall fold, their folding energetics is governed by the same quantitative pattern. Relative to the unfolded ensemble, the native conformation is enthalpically stabilized by comparable contributions from protein-protein and solvent-solvent interactions, and it is nearly equally destabilized by interactions between protein and solvent molecules. From the perspective of elementary physical interactions, the native conformation is stabilized by van de Waals and coulombic interactions and is destabilized by bonded interactions. Also common to the four proteins, the sign of the heat capacity change is set by protein-solvent interactions or, from the alternative perspective, by coulombic interactions.

## Introduction

Most proteins perform their biological functions once they have adopted a specific conformation among an enormity of alternatives available. Such special conformation is known as the native state and constitutes the most stable spatial arrangement polypeptide atoms can adopt within biologically relevant times when they are immersed, typically, in an aqueous solution^1^. Understanding the interactions between polypeptide and water molecules that drive the transformation of the initially unfolded polypeptide into its native folded conformation and determine its greater stability is a fundamental objective of Structural Biology^2^. From a practical point of view, achieving a detailed, quantitative understanding of the protein folding equilibrium may transform protein design into a trustable routine tool, which is much needed to perfect a growing range of biotechnological processes, including the fabrication of biological drugs^3^. Not of minor importance, it may contribute to boosting personalized medicine by setting the interpretation of genetic variants and their impact on human disease on firmer grounds^4^.

The stability of a protein native conformation is governed by the free energy difference (ΔG) of the equilibrium it establishes with the ensemble of unfolded conformations. While the experimental determination of ΔG is not usually complicated^5^, its interpretation in terms of specific atomic interactions^6,7^ remains a great challenge. Despite prolonged efforts by the biophysical community, the thermodynamic properties that determine protein forms cannot be calculated yet from first principles. Two such properties, the changes in enthalpy (ΔH) and in heat capacity (ΔCp) upon folding, can be measured directly from calorimetric experiments and are particularly amenable to interpretation^7–9^. As with purely biophysical approaches, protein engineering attempts to provide detailed knowledge of protein energetics at the molecular level have found difficulties in assigning differences in stability, or in the other thermodynamic magnitudes, to changes in specific elementary interactions (e.g. coulomb or van der Waals). This is because amino acid residue replacements characteristically summarize energy changes of several types^6,8,10^. On the other hand, Molecular Dynamics (MD) simulations have gained momentum in the study of protein energetics^11–13^. Improvements in force fields and water models, and continued increase in computation power have enabled a better sampling of the folding equilibrium, making atomistic MD simulation a well-suited approach to provide a precise dissection of protein folding energetics^14–17^.

At present, after intense scrutiny of protein energetics through experiment and simulation of natural proteins and designed variants, no consensus has emerged on the relative contributions of protein-protein, protein-solvent and solvent-solvent interactions to the changes in enthalpy and heat capacity that make native proteins stable at physiological temperatures^6,9,18,19^. The same uncertainty applies to the relative contribution of van der Waals and electrostatic interactions to those fundamental thermodynamic magnitudes^6,20^. To address this problem, we have recently developed a method^21,22^ that calculates ΔH and ΔCp of the protein folding/unfolding reaction from MD simulations of the native conformation and an ensemble of carefully generated unfolded ones^21–23^. This method, which circumvents the otherwise limited capability of MD approaches to simulate the protein folding time (from microseconds up to tens of seconds or beyond) as well as the structural overcompaction described for some classical MD force fields when combined with certain water models^15,24–27^, yields ΔH and ΔCp values in agreement with experiment^21,22^. Here, we provide a detailed dissection of four model proteins whose energetics have been successfully calculated using the indicated approach. The energy patterns derived from each of the four proteins coincide to reveal the long sought for signs and magnitudes of the chief contributions to protein folding energetics by the interacting molecules and elementary forces involved.

## Methods

### Proteins Simulated and Structural Models Used

Four proteins for which extensive reliable thermodynamic data of their folding/unfolding equilibrium are available have been simulated, namely: the ribonuclease from *Bacillus amiloliquefaciens* (barnase)^28^, the *Staphylococcus aureus* nuclease (SNase)^29^, the apoflavodoxin from *Anabaena* PCC 7119 (apoFld)^30^ and the chymotrypsin inhibitor 2 from barley (CI2)^31^. Their native ensembles have been represented by X-ray structures (PDB IDs: 1A2P^32^, 2SNS^33^, 1FTG^34^ and 2CI2^35^, respectively) with crystallographic resolutions of 1.5, 1.5, 2.0 and 2.0 Å, respectively. Their unfolded ensembles have been obtained by means of the ProtSA server^23^ that generates from an input sequence a large ensemble of unfolded conformations consistent with NMR and SAXS properties of fully unfolded proteins. Unfolded ensembles of over 2000 conformations have been generated for each of those proteins. To facilitate subsequent computation, the most elongated unfolded structures (around 10 %) have been discarded and 100 conformations of each model proteins have been randomly selected to represent their unfolded ensembles.

### MD Simulation Setup and Sampling of Folding Energetics

The analysis here presented relies on all-atom MD simulations reported recently^22^, wherein folded and unfolded structures were simulated in explicit water (Tip3p model^36^). A summary of the simulation setup and solvating conditions is given in **SI Table 1**. Charmm22 with CMAP correction version 2.0^37^ (Charmm, for simplicity) was used to simulate the proteins with Gromacs 2020 package^38^. The energetics of the folded state was obtained from averages of 40 replicas of each folded protein whereas that of the unfolded state from averages of 100 different unfolded conformations simulated. To avoid the compacting effect exerted by most force fields, the simulation approach relied on short (2-ns) trajectories, which ensures efficient sampling while avoids structural overcompaction^21,22^.

**Table 1.**
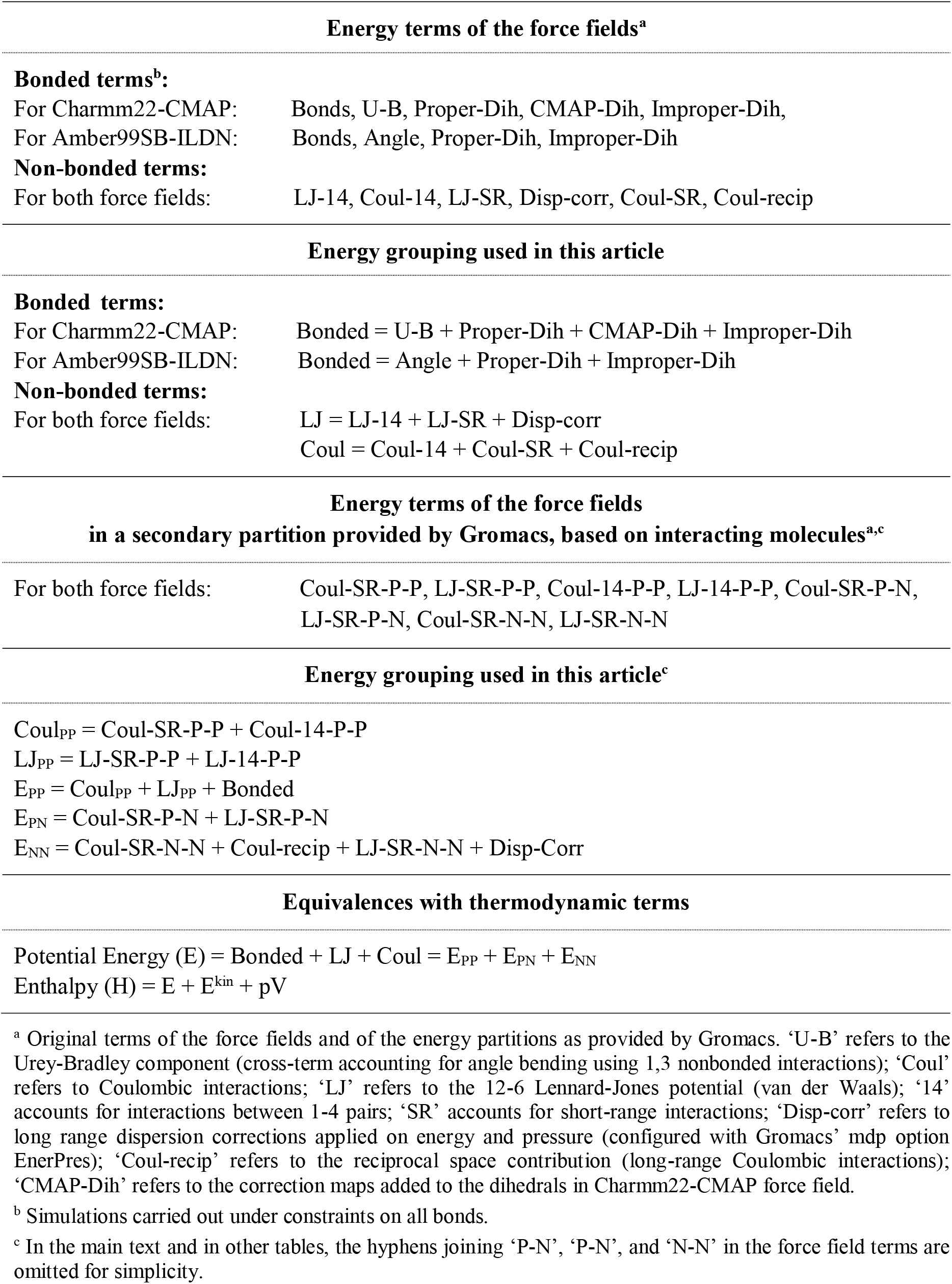
Energy inventories of the Charmm22-CMAP and Amber99SB-ILDN force fields, thermodynamic equivalences, and energy grouping used to discuss the contribution of specific elementary interactions (Bonded, Lennard-Jones or Coulomb) or specific molecular interactions: intraproteic (PP), protein-solvent (PN) or intrasolvent (NN) to protein folding ΔH and ΔC_p_

From the simulations of the folded and unfolded conformations of a given protein, ΔH of folding (ΔH_fol_) is calculated^21,22^ as the ensemble-averaged enthalpy of the simulated boxes containing folded protein minus that of the boxes containing unfolded conformations. For this ΔH_fol_ calculation to hold, boxes with folded protein must contain the same number of water molecules and counterions that the corresponding boxes with unfolded protein (**Figure 1**). For each protein, ΔH_fol_ is calculated at three temperatures and ΔCp_fol_ values are obtained as the slope of a representation of calculated ΔH_fol_ versus simulation temperature. Importantly, for the four proteins here considered, the calculated ΔH_fol_ and ΔCp_fol_ values agree with those determined experimentally, and the ΔG_fol_ values obtained from the calculated ΔH_fol_ and ΔCp_fol_ values plus the experimental melting temperature^39^ also agrees with the experimentally determined conformational stabilities^22^.

**Figure 1.**
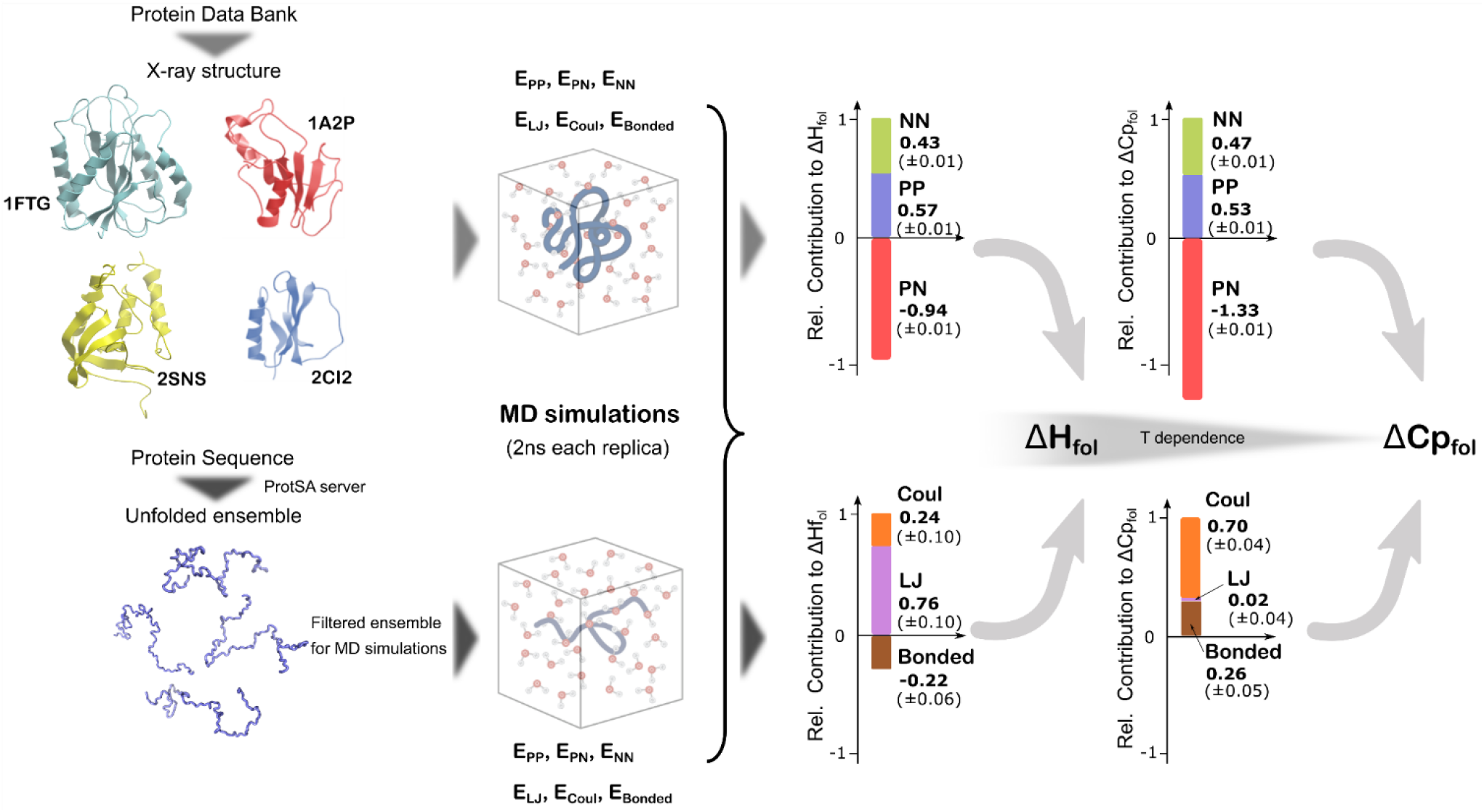
General scheme for the calculation of ΔH_fold_ and ΔCp_fold_ and their relative molecular and elementary contributions. The scheme part at the left-hand side -including the simulation boxes− indicates the simulated proteins and summarizes the MD setup followed, which is fully described in Galano-Frutos et. al.^22^. The bar graphs at the right-hand side show the averaged (standard errors between parentheses) molecular and elementary relative contributions to ΔH_fol_ and ΔCp_fol_ obtained for the four proteins analyzed. Positive relative contributions to ΔH_fol_ indicate molecular or elementary interactions that stabilize the folded conformation, while negative relative values indicate destabilizing contributions. In bars graphs showing relative contributions to ΔCp_fol_, averaged positive values indicate stabilizing contributions that become even more stabilizing with temperature, while averaged negative values indicate destabilizing contributions that become even more destabilizing with temperature. The reference value of 1 in the bars graphs represents the sum of the positive contributions. The negligible (slightly positive) van der Waals (LJ) average contribution to ΔCp_fol_ is the only contribution whose sign varies among the proteins analyzed.

Holonomic constraints on all the bonds have been applied through the MD simulations. A test aimed to assess the importance of the application of bond constraints in the analysis of the different energy terms evaluated here was performed for barnase by imposing constraints only on bonds involving hydrogen atoms. Besides, and for the sake of comparison, barnase energetics has been additionally calculated from simulations of folded and unfolded conformations using Amber99SB-ILDN^40^ (Amber for simplicity) combined with the same water model (Tip3p^36^) used with Charmm. In this case 10 replicas of the folded state and 40 unfolded structures were simulated^21^.

### Energy Partitions Provided by the Force Fields and Terminology Used for Grouped Terms

Charmm force field uses five energy terms to model the energy arising from the covalent structure, namely: bonds, angles with a Urey-Bradley correction, proper dihedral, CMAP-dihedral correction, and improper dihedral. In this work we will report the energies of all those terms grouped in a single energy term named ‘Bonded’ (**Table 1**). In our simulation approach, all bond distances have been restrained. Therefore, the value of the bonds energy term does not change with time nor it differs between the folded and unfolded conformations. To assess the influence of restraining bond energy on the values obtained for the other energy terms in the force field, we have compared barnase simulations wherein holonomic constraints have been applied to all bonds (**Table 2**) with barnase simulations where constraints have been only applied on bonds involving hydrogen atoms (**Table S2**). The comparison makes clear that, applying either of these two constraints configurations, the calculated contribution to folding enthalpy of the different bonded and non-bonded terms is not significantly altered (**Table 2** and **SI Table 2**).

**Table 2.**
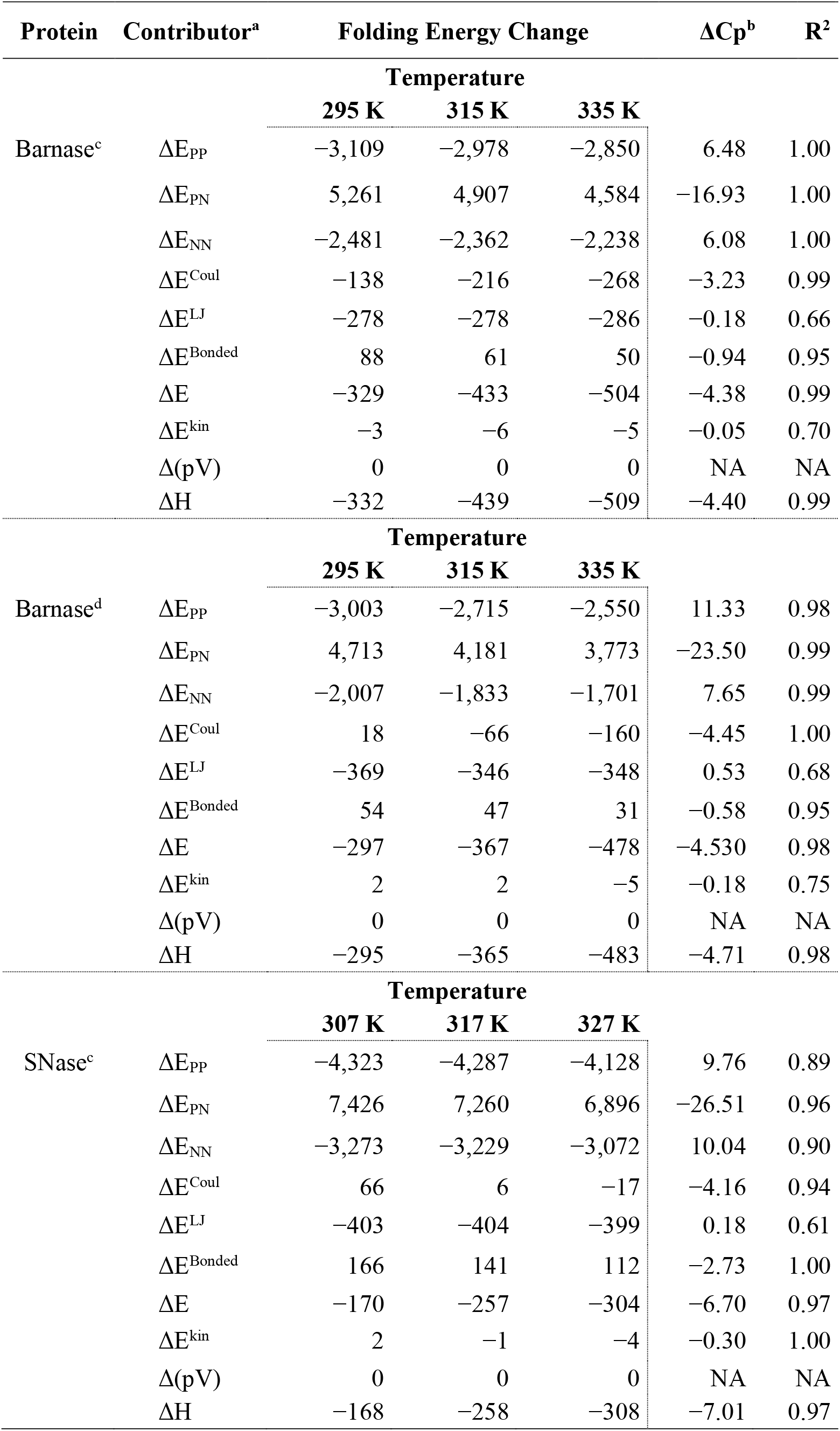
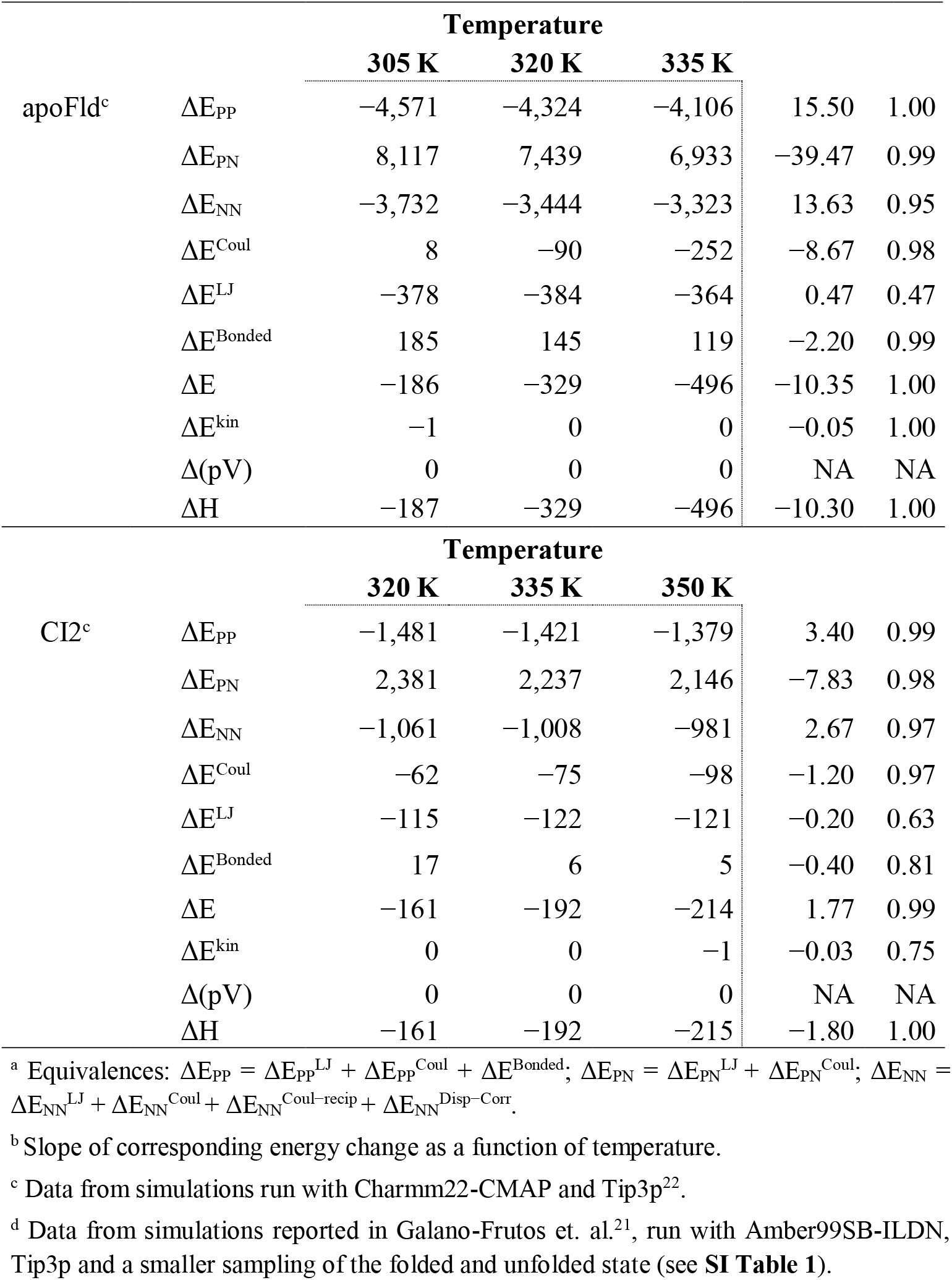
Contributions to ΔH and ΔCp of folding

Charmm models the non-bonded interactions established between protein atoms, between solvent molecules and between protein and solvent molecules by means of Lennard-Jones (LJ) and Coulomb (Coul) energy terms. Non-bonded interactions are computed distributed in six terms, namely, LJ-14, Coul-14, LJ-SR, Coul-SR, Disp-corr and Coul-recip (see detailed descriptions in **Table 1**). We will report them grouped in two terms: LJ (encompassing LJ-14, LJ-SR and Disp-corr) and Coulomb (encompassing Coul-14, Coul-SR and Coul-recip) (**Table 1**). In either force field, the time-averaged enthalpy (H) of a simulated trajectory containing protein and explicit water is obtained by summing up the Bonded, the LJ and Coul terms (together making up the potential energy, E), the kinetic energy (E^kin^) and the pV term (p = pressure, V = volume). The change in ΔH_fol_ can be then obtained as the enthalpy corresponding to the boxes containing folded protein (averaged over the simulated replicas) minus that of boxes containing unfolded protein (averaged in the same way)^21,22^, provided the number of water molecules and ions is the same in all boxes.

Alternatively, the energy terms obtained from the simulations can be grouped in a different manner to highlight the separate contribution of intraproteic (protein-protein), intrasolvent (solvent-solvent), and protein-solvent interactions. Indeed, GROMACS’s^38^ output provides the following terms: Coul-SR-PP, LJ-SR-PP, Coul-14-PP, LJ-14-PP, Coul-SR-PN, LJ-SR-PN, Coul-SR-NN, LJ-SR-NN, where PP refers to interactions between protein atoms, PN to interactions between protein and non-protein atoms (water molecules and ions, as the simulated proteins contain no cofactors), and NN to intrasolvent interactions. We will report the different PP terms grouped as Coul-PP (Coul-SR-PP + Coul-14-PP) and LJ-PP (LJ-SR-PP + LJ-14-PP). Thus, the potential energy due to intraproteic interactions (E_PP_) is obtained as Coul-PP + LJ-PP + Bonded, that corresponding to protein-solvent interactions (E_PN_) is Coul-SR-PN + LJ-SR-PN, and that related to solvent-solvent interactions (E_NN_) is, in principle, Coul-SR-NN + LJ-SR-NN (**Table 1**). However, two additional terms, Disp-corr and Coul-recip, also contribute to the potential energy of the simulated box. While these terms combine PP, PN and NN contributions, they essentially arise from solvent-solvent interactions, which represent >98 % of the pairwise interactions taking place in the simulated box. The contributions of the Disp-corr and Coul-recip terms have been thus added to those of Coul-SR-NN + LJ-SR-NN to jointly describe E_NN_. We would like to indicate that ignoring the Disp-corr and Coul-recip terms in our analysis would make very little difference, as their contribution to the change in solvent-solvent interaction energy upon folding is very small.

The Amber energy terms are quite similar to those in Charmm, which facilitates pair-wise comparisons of the values obtained for the same protein using either force field. The only difference is that Amber does not include CMAP dihedral correction, so that it uses four terms (bonds, angles, proper dihedral, and improper dihedral) to model the energy from the covalent structure. In the analysis of barnase simulations run with Amber, these four terms will be reported grouped in a single energy term named ‘Bonded’ (**Table 1**).

## Results

### Accuracy of Folding Enthalpy and Heat Capacity Changes Computed from Atomistic MD Simulations

In previous work^21,22^, we have shown that the changes in enthalpy and heat capacity associated to the folding/unfolding of several proteins can be accurately calculated from short atomistic MD simulations of the corresponding folded conformations and unfolded ensembles. The calculated enthalpy and heat capacity values (**Table 2**) correlate well with the experimental ones (**SI Table 3**). For the four proteins here analyzed, the correlation and slope of the linear plots are close to unity (for enthalpy changes: R^2^=0.99 and slope=0.98; for heat capacity changes: R^2^=0.99 and slope=0.86, **SI Figure 1**). We take this as a strong indication that the force field used here (Charmm22 with CMAP correction, version 2.0^37^) describes well the energetics of the folding equilibrium despite its known compacting effect on protein conformations^15,21,27^ that we simply skip by sampling the simulated folded and unfolded states through short 2-ns simulations^21,22^. In our experience, newer force fields specifically tuned to avoid compaction may not maintain the accuracy of protein enthalpy change calculations by difference^21^.

**Table 3.** Correlation coefficients (R^2^) between protein properties and different contributions to ΔH and ΔCp of folding^a^

| | Property | | | | Calculated contributions to $\Delta H$ | | | | | | $\Delta H^{\text{calc}}$ | $\Delta H^{\text{exp}}$ | Calculated contributions to $\Delta C_p$ | | | | | | $\Delta C_p^{\text{calc}}$ | $\Delta C_p^{\text{exp}}$ |
| --- | --- | --- | --- | --- | --- | --- | --- | --- | --- | --- | --- | --- | --- | --- | --- | --- | --- | --- | --- | --- |
|  |  |  |  |  | by partners |  |  | by force |  |  |  |  | by partners |  |  | by force |  |  |  |  |
| | Length | $\Delta \text{SASA}$ | $\Delta \text{SASA}_{\text{apol}}$ | $\Delta \text{SASA}_{\text{pol}}$ | PP | PN | NN | Bonded | LJ | Coul | | | PP | PN | NN | Bonded | LJ | Coul | | |
| Length | - | - | - | - | 0.97 | 0.99 | 0.98 | 0.96 | 0.94 | 0.04 | 0.16 | 0.06 | 0.90 | 0.94 | 0.97 | 0.83 | 0.83 | 0.80 | 0.95 | 0.97 |
| $\Delta \text{SASA}$ | 0.99 | - | - | - | 0.92 | 0.93 | 0.94 | 0.93 | 0.87 | 0.03 | 0.15 | 0.06 | 0.96 | 0.98 | 0.99 | 0.76 | 0.88 | 0.89 | 0.99 | 0.99 |
| $\Delta \text{SASA}_{\text{apol}}$ | 0.94 | 0.98 | - | - | 0.84 | 0.86 | 0.88 | 0.85 | 0.78 | 0.01 | 0.17 | 0.08 | 0.99 | 0.99 | 0.98 | 0.63 | 0.89 | 0.95 | 0.99 | 0.98 |
| $\Delta \text{SASA}_{\text{pol}}$ | 0.96 | 0.91 | 0.82 | - | 0.99 | 0.99 | 0.97 | 0.99 | 0.97 | 0.10 | 0.10 | 0.02 | 0.77 | 0.83 | 0.89 | 0.94 | 0.75 | 0.64 | 0.85 | 0.91 |

<sup>a</sup> Analysis carried out with calculated thermodynamics from simulations of barnase, SNase, apoFld and CI2. Correlation coefficients obtained from linear correlation with properties in the first column of the table.

All the energy differences contributing to ΔH_fol_ that are defined in **Table 1** have been calculated as time and replica average differences between enthalpy data pertaining to simulation boxes containing folded conformations and enthalpy data from boxes containing unfolded conformations. The primary “folded” and “unfolded” data corresponding to the four proteins here analyzed are shown in **SI Tables 3-7**.

### Contribution of Intraproteic and Solvation Interactions to ΔH_fol_

The protein folding enthalpy change is a consequence of the rearrangement of interactions between protein and solvent. The relative contribution to ΔH_fol_ of changes in intraproteic (ΔE_PP_), intrasolvent (ΔE_NN_) and protein-solvent (ΔE_PN_) interactions has been determined for barnase using the Charmm22-CMAP force field (**Table 2**). According to Charmm, the enthalpic stabilization of the barnase native conformation at 42 °C (−438 kJ/mol) is driven by a slightly larger stabilizing contribution of intraproteic interactions (ΔE_PP_ = −2978 kJ/mol) which is opposed by the destabilizing combination of the two solvation contributions (ΔE_PN_ + ΔE_NN_ = +2545 kJ/mol). An additional minor contribution of −6 kJ/mol from the kinetic energy change completes the enthalpy balance. This contribution likely arises from a non-perfect cancelation of the theoretically identical but very large kinetic energies of the folded and unfolded simulation boxes. The indicated overall destabilizing effect of solvation contributions is composed of two opposing terms: a dominant, destabilizing term due to protein-solvent interactions (ΔE_PN_ = +4907 kJ/mol) and a stabilizing one due to intrasolvent interactions (ΔE_NN_ = −2362 kJ/mol, **Figure 2a**). Therefore, as barnase folds establishing new internal stabilizing interactions and loosing preexisting stabilizing interactions with solvent to a much larger extent, new interactions are also formed among surrounding solvent molecules, which makes an important stabilizing contribution to the native conformation. This stabilizing contribution is almost as large as that of intraproteic interactions.

**Figure 2.**
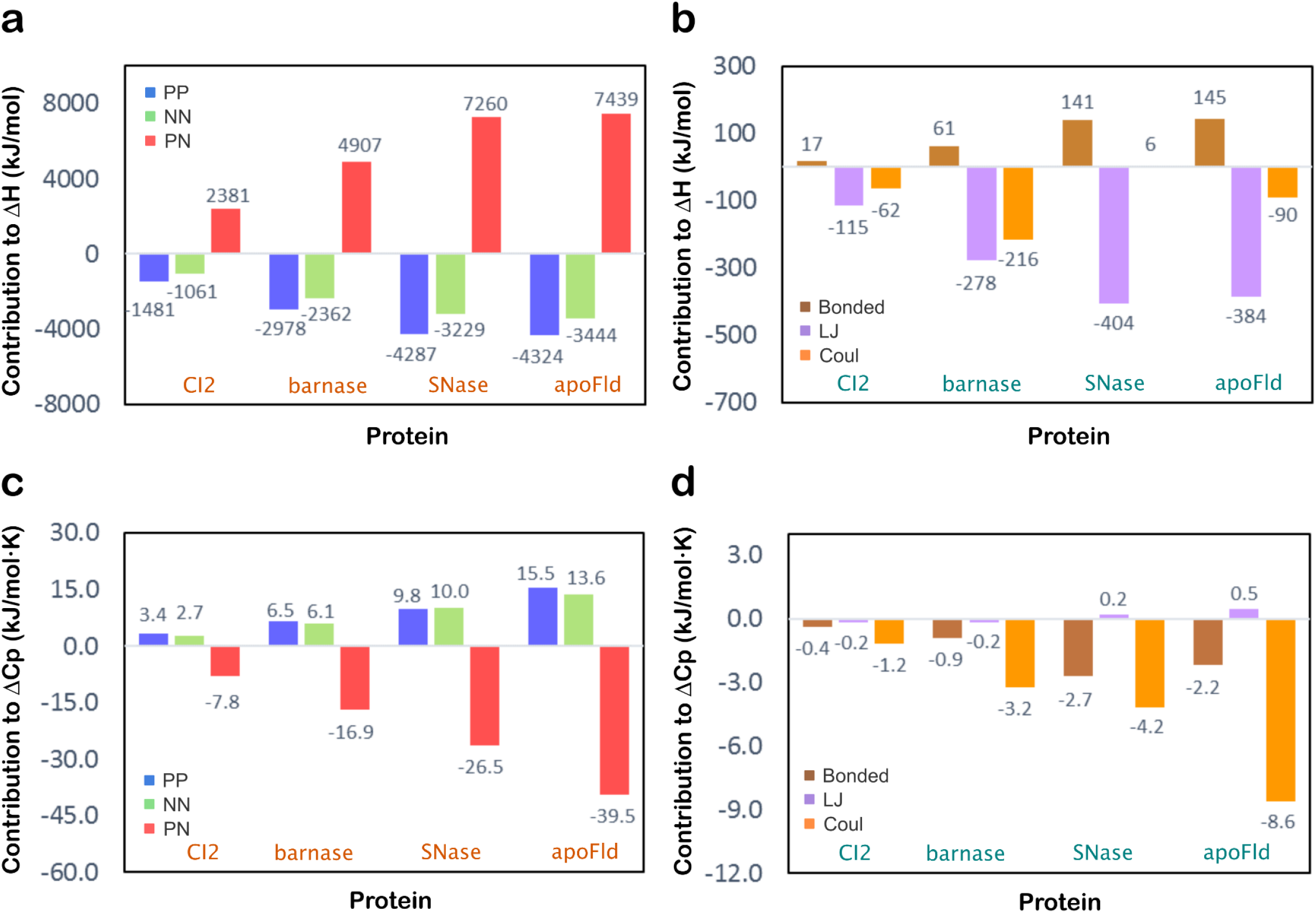
Contributions of molecular interactions and of elementary interactions to ΔH_fol_ (a, b) and ΔCp_fol_ (c, d). ‘PP’, ‘NN’ and ‘PN‘ in panels **a** and **c** refer to intraproteic, intrasolvent and protein-solvent energy contributions, respectively, whereas ‘Bonded’, ‘LJ’ and ‘Coul’ and in panels **b** and **d** refers to bonded, Lennard-Jones (van der Waals) and coulombic (electrostatics) contributions, respectively.

For the sake of comparison and to discard that the folding pattern provided by Charmm could be force field specific rather than intrinsic to the protein simulated, we have performed the same energy dissection of barnase folding using data obtained from simulations performed with Amber99SB-ILDN in otherwise identical conditions. Clearly, Amber captures the same pattern. According to Amber, the contribution of intraproteic interactions is −2715 kJ/mol and those of the two solvation terms are +4181 and −1833 kJ/mol, respectively (**Table 2**). Thus, two independent force fields, Charmm and Amber, coincide in revealing an important stabilizing contribution of intrasolvent interactions.

To assess whether the above pattern describing barnase folding energetics generally accounts for the relative contribution of intraproteic, protein-solvent and intrasolvent interactions to the folding enthalpy change of globular proteins, MD simulations have been carried out in the same temperature range for SNase (at 44 °C), apoFld (at 47 °C) and CI2 (at 47 °C) using Charmm (**Table 2**). Indeed, the same pattern is found in these three additional proteins, the factor describing the relative stabilizing contribution of intrasolvent and intraproteic interactions to ΔH_fol_ being 0.75 for SNase, 0.80 for apoFld and 0.72 for CI2, all of them very similar to the factor of 0.79 previously found for barnase. Moreover, a large cancellation of those stabilizing contributions is observed in the four proteins, which comes from destabilizing protein-solvent interactions. Namely, protein-solvent destabilizing interactions (ΔE_PN_) are, in absolute value, 0.92 (barnase), 0.97 (SNase), 0.96 (apoFld) or 0.94 (CI2) times as high as the combined stabilizing interactions, i.e. ΔE_PP_ + ΔE_NN_, of the corresponding protein (**SI Figure 2a**).

### Contribution of Bonded, Coulomb and Lennard-Jones interactions to ΔH_fol_

The folding enthalpy change of a protein can be partitioned differently attending to the elementary forces governing the reaction, rather to the participating molecules. At the indicated temperature of 42 °C, and under the Charmm force field, the barnase enthalpy change (ΔH_fol_ = −438 kJ/mol) arises from stabilizing energy contributions of coulombic (ΔE^Coul^ = −216 kJ/mol) and Lennard-Jones (ΔE^LJ^ = −278 kJ/mol) interactions, which are partly opposed by moderately destabilizing bonded interactions (ΔE^Bonded^ = +61 kJ/mol, **Table 2** and **Figure 2b**). It seems that, as barnase folds, LJ interactions −and to a lower extent coulombic interactions− strengthen at the expense of introducing some strain in the native conformation. The Amber force field reveals the same pattern for barnase folding as it also reports stabilizing contributions from coulombic and LJ interactions of −66 and −346 kJ/mol, respectively, and a moderate destabilizing contribution from bonded ones of +47 kJ/mol (**Table 2**). In agreement with this, analysis of the folding pathway of peptides or mini proteins (from 10 to 35 residues) using other force fields of the Amber family^41^ also showed a destabilizing contribution from bonded interactions.

To assess whether this is a general pattern characteristic of the protein folding reaction of globular proteins, the ΔH_fol_ values calculated for SNase, apoFld and CI2 at similar temperatures using Charmm (see above) has also been partitioned into coulombic, LJ and bonded contributions. As in the case of barnase, the major stabilization for all these proteins comes from LJ interactions and there is a clear destabilizing contribution from bonded ones. The contribution of coulombic interactions to ΔH_fol_ is more variable in the four proteins, as it is clearly stabilizing in barnase, apoFld and CI2 but it is close to negligible for SNase at the temperature of comparison (**Table 2** and **Figure 2b**).

### Temperature Dependence of Intraproteic and Solvation Interactions: contributions to ΔCp_fol_

The folding enthalpies (ΔH_fol_) of barnase, SNase, apoFld and CI2 have been determined at three temperatures using the Charmm force field and their corresponding ΔE_PP_, ΔE_PN_ and ΔE_NN_ components have been obtained at each temperature. Then, their individual contributions to the heat capacity of folding (ΔCp_fol_) have been calculated as the slopes of the corresponding linear plots of ΔE terms versus simulation temperature. These plots have shown excellent correlation coefficients (R^2^) of 1.00 for barnase, 0.88-0.96 for SNase, 0.95-1.00 for apoFld and 0.97-0.99 for CI2 (**Table 2**). In barnase, the contribution of intraproteic interactions (ΔCp_PP_) to ΔCp_fol_ is +6.48 kJ/mol∙K (**Table 2**), indicating that intraproteic interactions reduce their stabilizing effect on the folded conformation as the temperature is raised. On the other hand, the overall contribution of the solvation terms (ΔCp_PN_ + ΔCp_NN_) is −10.85 kJ/mol∙K (**Table 2**), which means that solvation destabilizes the folded conformation less and less as the temperature raises. As this decrease in solvation destabilization with temperature is more steep than the concomitant decrease of intraproteic stabilization, the overall enthalpy change stabilizes the folded state more as the temperature rises, which is reflected in the calculated ΔCp_fol_ of −4.40 kJ/mol∙K (**Table 2**). The two solvation terms make opposite contributions to this global negative value of ΔCp_fol_, namely, that of protein-solvent interactions is negative (ΔCp_PN_ = −16.93 kJ/mol∙K), while that of intrasolvent interactions is positive (ΔCp_NN_ = +6.08 kJ/mol∙K), **Table 2** and **Figure 2c**). Thus, as the temperature rises, the two stabilizing contributions of the native conformation (intraproteic and intrasolvent interactions) are reduced but the destabilizing protein-solvent interactions are reduced to a greater extent. The combined effect makes the enthalpic stabilization of the folded state increase with temperature. This pattern is also revealed when barnase is simulated using the Amber force field (**Table 2**). Importantly, analysis of the SNase, apoFld and CI2 simulations performed with Charmm (**Table 2**) indicates that the partition of their ΔCp_fol_ values into contributions from intraproteic, protein-solvent and intrasolvent interactions is basically the same as that described for barnase (**Figure 2c**). In the four proteins, positive and similarly large intraproteic and intrasolvent contributions appear counterbalanced by a negative, larger contribution of protein-solvent interactions, which exceeds the positive contributions by a factor ranging from 1.29 to 1.36 (**SI Figure 2b**).

### Temperature Dependence of Bonded, Coulomb and Lennard-Jones Interactions: Contributions to ΔCp_fol_

The calculated ΔCp_fol_ values of the four proteins can also be partitioned into specific contributions associated to elementary interactions: LJ, Coulomb and bonded. For the four proteins, the energy versus temperature linear plots of those contributions allow us to calculate the coulombic and bonded contributions to ΔCp_fol_, which show high correlation coefficients (R^2^) ranging from 0.94 to 0.99 for coulombic, and from 0.81 to 1.00 for bonded. In contrast, the LJ contribution to ΔCp_fol_ is calculated with lower R^2^ coefficients (from 0.47 to 0.81). This LJ contribution to ΔCp_fol_ is consistently small for the four proteins (−0.18, +0.20, +0.47 and −0.20 kJ/mol∙K), which indicates that the LJ balance of their folding reactions is rather insensitive to temperature. The ΔCp_fol_ associated to coulombic interactions makes by far the larger contribution in the four proteins, and there is also a significant contribution from bonded interactions in all cases (**Figure 2d**). The coulombic and bonded contributions to ΔCp_fol_ are both negative, which indicates that the stabilizing effect of coulombic interactions increases with temperature while the destabilizing effect of bonded interactions decreases. As the strong stabilizing effect of the native conformation exerted by LJ interactions appears to be rather temperature insensitive, the larger enthalpic stabilization of the native conformation at high temperatures compared to lower temperatures does not arise from LJ interactions. It arises from a strengthening of stabilizing coulombic interactions combined with a reduction of destabilizing bonded ones.

### Correlation of ΔH_fol_ with Protein Length and Changes in Solvent Exposure

There has been significant efforts to correlate folding enthalpy and heat capacity changes to simple properties of polypeptides such as protein size or solvent exposure^9,19^. Our analyses of the barnase, SNase, apoFld and CI2 MD simulations (**Table 3**) show (**Figure 3a**) that correlations between individual ΔE_PP_, ΔE_PN_ and ΔE_NN_ contributions to ΔH_fol_ and protein length (**SI Table 8**) are impressive (R^2^=0.97-0.99), yet the correlation of ΔE (which combines the three energy terms) with protein length is poor (R^2^=0.16). This is so because, for each protein, the individual ΔE_PP_, ΔE_PN_ and ΔE_NN_ contributions are one order of magnitude higher than their added value (ΔE), and small errors in their calculated values can translate into a big error in ΔE. As ΔE only differs from ΔH_fol_ by a rather small kinetic energy term, it follows that the correlation observed between calculated ΔH_fol_ and protein length is similarly poor (R^2^=0.16, **Figure 3e**). This agrees with the fact that the experimental ΔH_fol_ values for these proteins (**SI Table 8**) do not correlate with protein length any better (R^2^=0.06, **Figure 3e**). Considering the alternative partition of enthalpy changes attending to elementary forces, both stabilizing LJ and destabilizing bonded interactions correlate with protein length (R^2^=0.94 and 0.96, respectively) but coulombic interactions do not (R^2^=0.04, **Figure 3c**), which also explains from this angle the poor correlation of calculated ΔH_fol_ with protein length (R^2^=0.16, **Figure 3e**).

**Figure 3.**
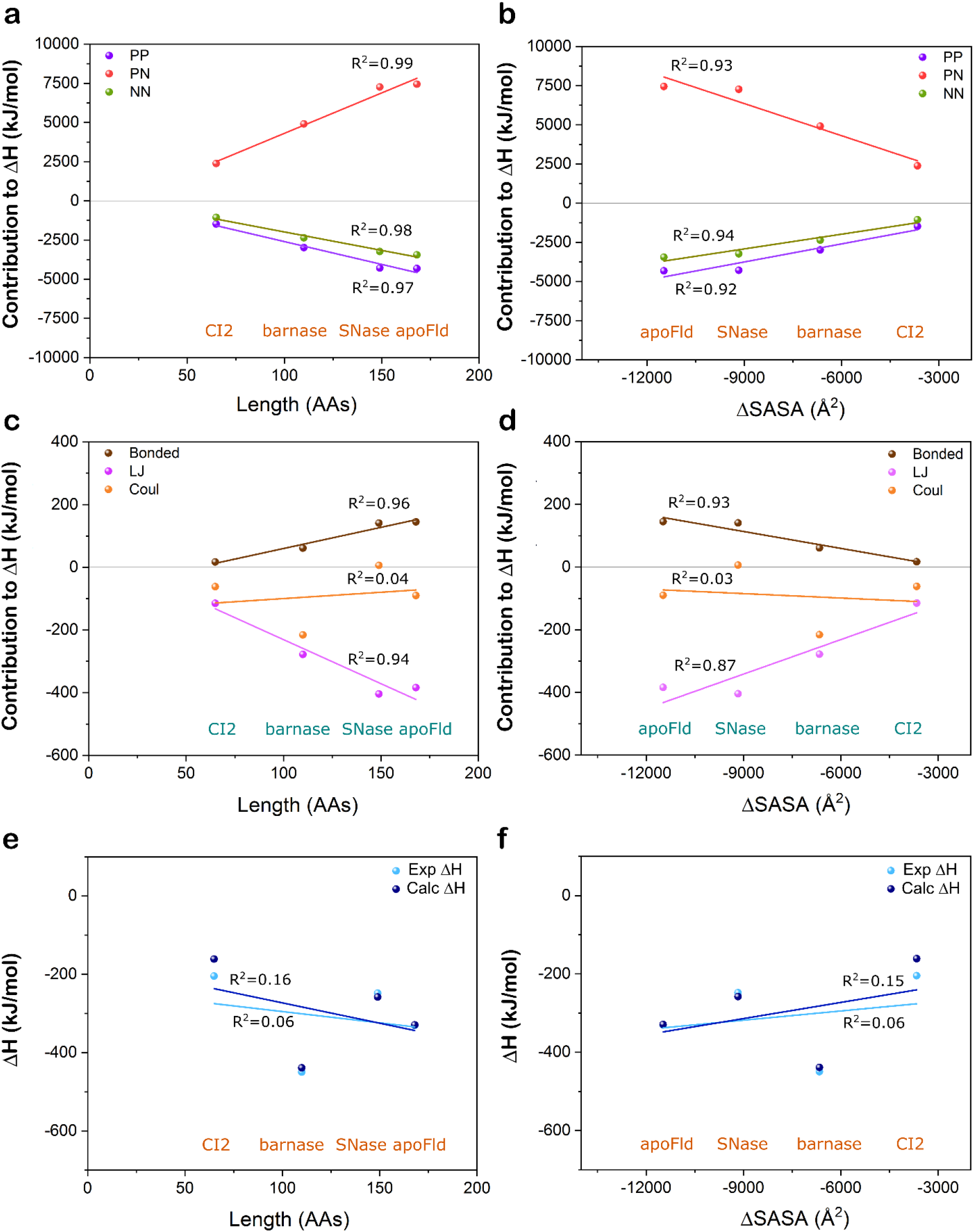
Linear correlation between molecular (a, b) and elementary (c, d) contributions to ΔH_fol_ and protein length (a, c) or ΔSASA (b, d). Linear correlation between calculated and experimentally determined ΔH_fol_ and protein length (number of amino acid residues) (e) or ΔSASA (d). ‘PP’, ‘NN’ and ‘PN’ in **a** and **b** refer to intraproteic, intrasolvent and protein-solvent energy contributions, respectively, whereas ‘Bonded’, ‘LJ’ and ‘Coul’ in **c** and **d** refer to bonded, Lennard-Jones (van der Waals) and coulombic (electrostatics) contributions, respectively. Squared Pearson correlation coefficients are included close to each fitting line.

On the other hand, the folding change in SASA (ΔSASA), and its polar and apolar components (ΔSASA_pol_ and ΔSASA_apol_) have been calculated for each of these proteins (**SI Table 8**)^23^. Individual ΔE_PP_, ΔE_PN_ and ΔE_NN_ energy contributions to ΔE correlate best (**Table 3**) with ΔSASA_pol_ (R^2^ from 0.97 to 0.99), then with ΔSASA (R^2^ from 0.92 to 0.94, **Figure 3b**), and then with ΔSASA_apol_ (R^2^ from 0.84 to 0.88). However, as seen above for the correlations with protein length and likely for the same reason, ΔE does not correlate with either total, polar or apolar changes in SASA (R^2^ from 0.11 to 0.18). Consequently, calculated ΔH_fol_ values do not correlate either with any of the SASA changes (R^2^ from 0.10 to 0.17) (**Table 3** and **Figure 3f**). This agrees with the lack of correlation between SASA changes and experimental ΔH_fol_ values (R^2^ from 0.02 to 0.08). For the alternative partition of enthalpy changes into elementary contributions, both the bonded and LJ terms correlate either strongly or moderately with ΔSASA (R^2^=0.93 and 0.87, respectively, **Figure 3d**) and its polar and apolar components (R^2^ values from 0.78 to 0.99, **Table 3**). In contrast, the Coulomb term does not correlate with the different ΔSASAs (R^2^ from 0.01 to 0.10, **Table 3** and **Figure 3d**).

The very high correlations found between protein length or changes in SASA and the individual energy terms (ΔE_PP_, ΔE_PN_ and ΔE_NN_) totaling ΔE invite to calculate ΔH_fol_ from simple linear relationships. Unfortunately, as explained above, once those individual terms are summed, the resulting ΔE values no longer correlate with either length or SASA changes. For the alternative energy partition, while LJ and bonded interactions correlate well with protein length and with the various changes in SASA considered, coulombic interactions do not, which also explains the lack of correlation here observed between protein length or SASA changes and ΔH_fol_. In practice, no accurate calculation of ΔH_fol_ from protein length or from changes in SASA seems possible.

### Correlation of ΔCp_fol_ with Protein Length and Changes in Solvent Exposure

The correlation between ΔCp_fol_ (and molecular or elementary contributions to it) and protein length has also been examined. Intraproteic (ΔCp_PP_), protein-solvent (ΔCp_PN_) or intrasolvent (ΔCp_NN_) contributions to ΔCp_fol_ correlate well (R^2^ from 0.90 to 0.97) with protein length (**Table 3** and **Figure 4a**), and the combined calculated ΔCp_fol_ also do so (R^2^=0.95, **Table 3** and **Figure 4e**). In agreement with this, the correlation of experimental ΔCp_fol_ values with protein length is also high (R^2^=0.97, **Table 3** and **Figure 4e**). Considering the alternative energy partition, the contribution of bonded, LJ and coulombic interactions to ΔCp_fol_ all correlate moderately well with protein length (R^2^ from 0.80 to 0.83, **Table 3** and **Figure 4c**).

**Figure 4.**
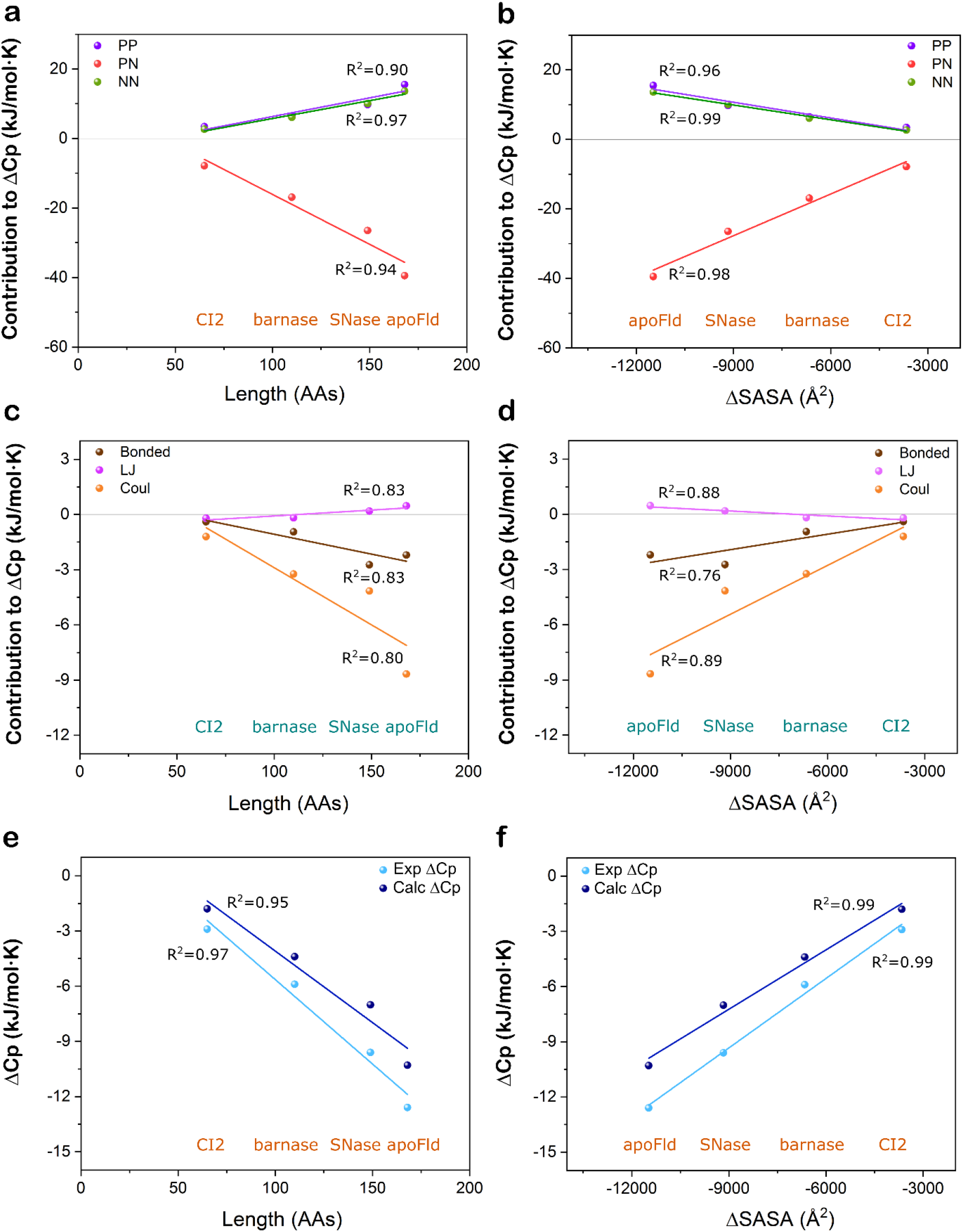
Linear correlation between molecular (a, b) and elementary (c, d) contributions to ΔCp_fol_ and protein length (a, c) or ΔSASA (b, d). Linear correlation between calculated and experimentally determined ΔCp_fol_ and protein length (number of amino acids) (e) or ΔSASA (d). ‘PP’, ‘NN’ and ‘PN’ in **a** and **b** refer to intraproteic, intrasolvent and protein-solvent energy contributions, respectively, whereas ‘Bonded’, ‘LJ’, and ‘Coul’ in **c** and **d** refers to bonded, Lennard-Jones (van der Waals) and coulombic (electrostatics) contributions, respectively. Squared Pearson correlation coefficients are included close to each fitting line.

On the other hand, ΔCp_PP_, ΔCp_PN_ and ΔCp_NN_ contributions to ΔCp_fol_ show high correlations with total ΔSASA (R^2^ from 0.96 to 0.99, **Table 3** and **Figure 4a**), even better correlation with ΔSASA_apol_ (R^2^ from 0.98 to 0.99, **Table 3** and **SI Figure 3a**), and a clear but lower correlation with ΔSASA_pol_ (R^2^ from 0.77 to 0.89, **Table 3** and **SI Figure 3c**). For the alternative partition, the three contributions (ΔCp_Bonded_, ΔCp_LJ_ and ΔCp_Coul_) correlate (R^2^ from 0.63 to 0.95, **Table 3**) with total ΔSASA (**Figure 4e**) and apolar (**SI Figure 3b**) or polar SASA changes (**SI Figure 3d**). While the LJ contribution correlates similarly well with polar and apolar changes, the coulombic one correlates better with ΔSASA_apol_ and the bonded contribution with ΔSASA_pol_. Overall, the calculated ΔCp_fol_ correlates best with either ΔSASA or its apolar component (R^2^=0.99) and the experimental ΔCp_fol_ is also highly correlated with either of these two changes (R^2^=0.98-0.99).

In practice, ΔCp_fol_ can be calculated accurately, at least for these proteins, with either of these linear equations: ΔCp_fol_ = 3.53 − 0.0917 × Length (R^2^=0.97); ΔCp_fol_ = 1252 − 73.1 × ΔSASA (R^2^=0.99), where Length is the number or amino acid residues in the protein and ΔSASA is the change in total SASA as calculated using ProtSA^23^.

## Discussion

### The Difficulty of Dissecting Protein Folding Energetics from Experiments

Some basic facts of the protein folding equilibrium that have been known for a long while are: 1) proteins that fold into compact native conformations tend to be stable at physiological temperatures and to unfold as temperature increases; 2) relative to unfolded ensembles, native conformations are stabilized at physiological temperatures by enthalpy and destabilized by entropy; 3) enthalpy differences are due to a combination of intraproteic interactions, protein-solvent interactions and interactions between solvent molecules; 4) the main interactions established among protein and solvent atoms are driven by electrostatic and van der Waals forces. However, despite this consolidated knowledge, the relative contribution of the different players to the stabilization of the native conformation is controversial, since a myriad of concurrent interactions take place simultaneously in the protein folding equilibrium that cannot be measured independently.

Before the advent and generalization of protein engineering, extensive efforts were done to model protein energetics based on thermodynamic data available for polar and apolar model compounds, analyses of protein composition, and estimations of the changes in SASA upon folding as a proxy for changes taking place in protein solvent interactions^42–47^. These studies were fostered by the development of precise calorimeters and led to proposals attributing a major contribution to ΔCp_fol_ to hydration interactions. The rational for this proposal was that, in the unfolded state, the interactions between solvent molecules and apolar groups are more intense than in the folded one, which explains the sign of ΔCp_fol_^9,18,19,48,49^. However, in spite of extensive analysis and modelling, the relative contributions to ΔCp_fol_ of the two solvation terms involved (due to protein-solvent and intrasolvent interactions) and of protein-protein interactions has remained an open question^18^. Protein engineering has tried to clarify this problem by comparing the energetics of very similar protein variants usually differing in one amino acid residue. This approach has been hampered by limitations, which still exist^50,51^, on the chemical changes that could be engineered and, more fundamentally, by the fact that replacement of specific chemical groups often affects more than one type of interaction, as it has been discussed^6^. Besides, protein engineering has predominantly focused on determining free energy changes rather than enthalpy or heat capacity changes, which is not ideal for the purpose of performing a fine dissection of protein energetics. Altogether, despite the large number of mutational experiments done, which keep feeding databases^52,53^ used to train protein stability predictors^54,55^, the relative contribution of the different elementary interactions to protein stability remains unclear.

More recently, atomistic simulation of proteins has opened a new window into detail. MD simulations, the most popular and suited simulation approach to study protein stability and folding^6,56–59^, enables computing protein and solvent energetics using potential energy functions, force fields, whose individual terms are related to specific forces. It also uses integration procedures that can classify the interactions as involving only protein atoms, only solvent atoms or as taking place between atoms of both protein and solvent molecules. The force fields typically used in atomistic MD simulations contain multiple parameters primarily derived from experimental data or from quantum chemical calculations often obtained from small molecules^60^. Although it may be feared that this reductionist approach may lead to insufficiently accurate calculation of the properties of complex molecules, such as proteins, the approach has a positive side. Since the data used in the force field parameterization are not primarily obtained from the large simulated molecules (i.e. proteins), the values of properties calculated for them will not be merely returning information previously incorporated into the force field. We have recently shown that, using atomistic MD simulations, the changes in enthalpy that take place when a polypeptide that is dissolved in water folds to its native conformation can be calculated accurately^21,22^. In this approach, simulations of the folded structure and of a representative sample of the unfolded ensemble are performed and the enthalpy changes are obtained by subtracting the ensemble-averaged enthalpies of such states^21,22^. In addition, the temperature dependence of the enthalpy change, i.e. the heat capacity change of folding/unfolding, can also be calculated within experimental error. Accurate calculation of ΔH_fol_ and ΔCp_fol_ from MD simulations has opened the possibility to dissect −beyond experiment− the contribution of elementary forces to protein folding/unfolding energetics and to clarify the specific role played by protein and solvent inter and intramolecular interactions. A coherent view of protein folding energetics emerges from dissection of the different contributions to ΔH_fol_ in four unrelated proteins whose MD-calculated and experimentally-determined ΔH_fol_ values coincide within error (**SI Figure 1**)^21,22^.

### A Coherent View of Protein Folding Energetics from MD Simulation

From the molecular perspective, folding enthalpy changes summarize rearrangements of internal protein (ΔE_PP_) and internal solvent (ΔE_NN_) interactions, as well as rearrangement of the interactions established between protein and solvent molecules (ΔE_PN_). In the four proteins here dissected, at similar temperatures around 45 °C, the folding reaction takes place amid a strong cancelation of stabilizing and destabilizing contributions that can be quantitatively described by equations 1 and 2. The native conformation is stabilized by the strengthening of both protein internal and solvent internal interactions, their relative contributions to ΔH_fol_ being similar and rather constant:

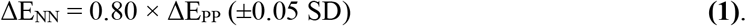

Conversely, the native conformation is intensely destabilized by a reduction of protein-solvent interactions, which is almost as intense as the combination of the stabilizing contributions:

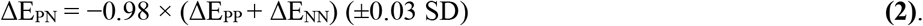

In other words, as the polypeptide folds (**Figure 5a**), the interactions it forms internally do not compensate the interactions it loses with the solvent molecules. However, stabilizing interactions concomitantly formed among solvent molecules drive the overall balance in favor of the native conformation. In terms of energy, almost half of the protein-solvent interactions that are lost when the protein folds are compensated by the new interactions that are established between solvent molecules.

**Figure 5.**
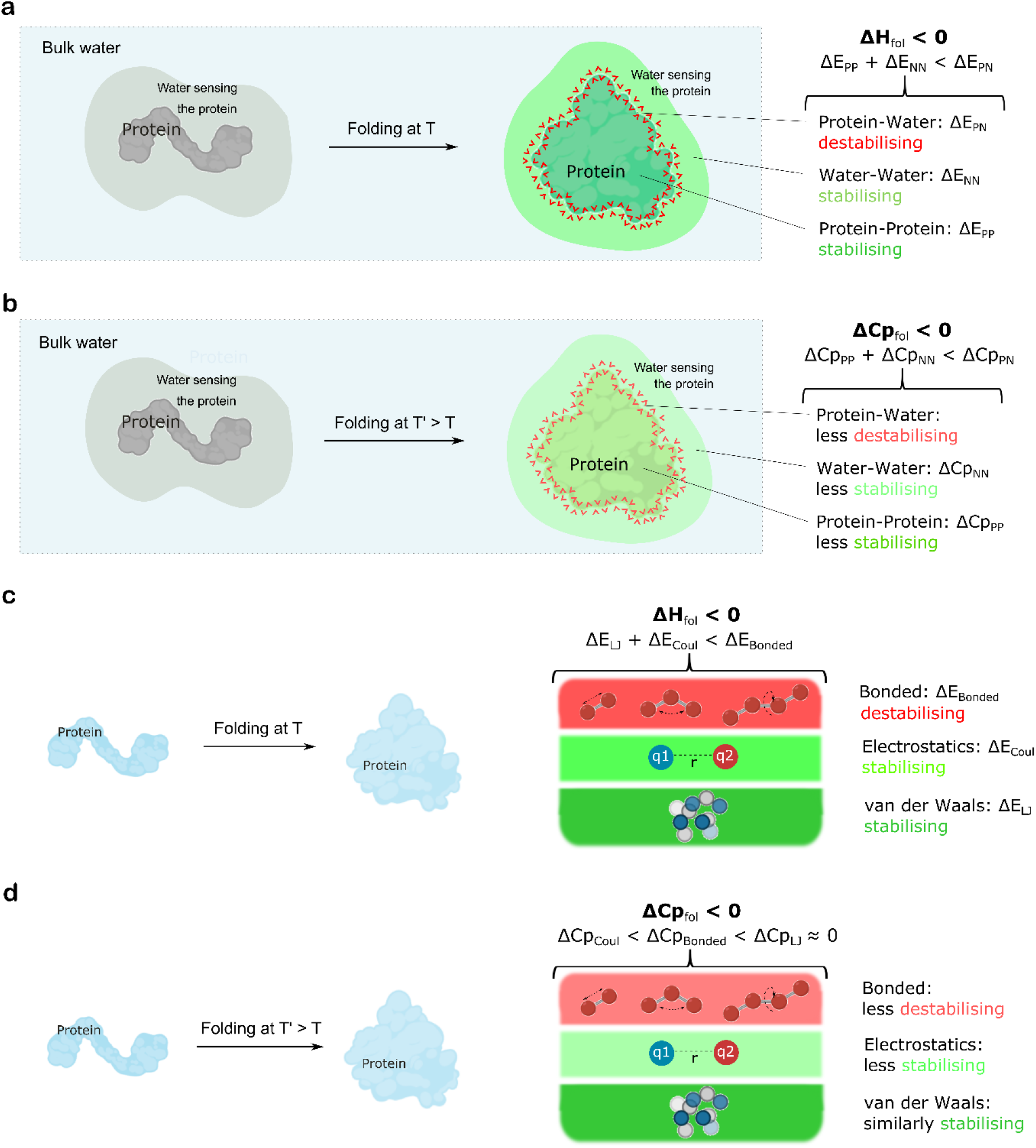
Contributions of molecular interactions and elementary forces to ΔH_fol_ and ΔCp_fol_. Panels **a** and **b** illustrate the energetics of protein folding and its temperature dependence, focusing on inter and intramolecular interactions between protein and solvent (essentially water) atoms. The unfolded system at the left (protein and non-bulk surrounding water molecules sensing the protein) is taken as the energy reference and is depicted in grey. The folded system at the right is colored differently to distinguish interactions that stabilize (water-water and protein-protein interactions; green shades) or destabilize (protein-water interactions, red lines) the system and therefore the folded conformation relative to the unfolded one. The temperature effect on the different molecular interactions shaping ΔH_fol_ determines how much they contribute to ΔCp_fol_. The temperature effects can be understood by comparing the comments at the right hand side of panels **a** (lower temperature) and **b** (higher temperature). Relative values of the different molecular contributions to ΔH_fol_ and ΔCp_fol_ and indicated qualitatively. Panels **c** and **d** illustrate the energetics of protein folding, focusing on elementary (bonded, van der Waals and electrostatic) interactions without differentiating between protein and solvent atoms. In these panels, the interactions stabilizing the folded state (van der Waals and electrostatic) are represented in green strips while the destabilizing contribution of bonded interactions arising from the protein covalent structure is indicated in the red strip on top of the green ones. The contribution of the different elementary interactions to ΔCp_fol_ can be understood by comparing the comments at the right hand side of panels **c** (lower temperature) and **d** (higher temperature). Relative values of the different elementary contributions to ΔH_fol_ and ΔCp_fol_ and indicated qualitatively.

From the perspective of elementary forces, a clear pattern, in this case qualitative, is also evident: i) the major stabilizing contribution to ΔH_fol_ is done by van der Waals interactions; ii) coulombic interactions also contribute to stabilize the native conformations but to a lesser extent; iii) stabilizing van der Waals and coulombic contributions are moderately opposed by destabilizing bonded contributions. As it appears (**Figure 5c**), protein folding in water is driven by a strengthening of van der Waals and coulombic interactions, at the cost of introducing some strain in the folded polypeptide. In agreement with this, avoiding local backbone strain has been reported to be critical for success in protein design^61^.

### Why ΔCp_fol_ is Negative?

As dictated by the negative sign of ΔCp_fol_^18^, raising temperature increases the enthalpic stabilization of proteins. From a molecular perspective, our analysis reveals a consistent cancellation of opposing contributions to ΔCp_fol_. On one hand, the contributions of both intraproteic and intrasolvent interactions (ΔCp_pp_ and ΔCp_NN_) are positive, meaning that, as temperature increases, both ΔE_PP_ and ΔE_NN_ stabilize the native conformation less and less (**Figure 5b**). Their relative impact on ΔCp_fol_ can be quantitatively described as:

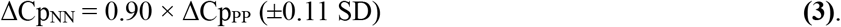

On the other hand, this weakening of the folded conformation as temperature increases is more than compensated by the negative and larger contribution of ΔCpPN, reflecting that the debilitation of the folded state by protein-solvent interactions, ΔE_PN_, is also markedly reduced at higher temperatures. As it appears (**Figure 5b**), increasing temperature weakens both the stabilizing and destabilizing molecular contributions to ΔH_fol_ but it weakens more steeply the destabilizing protein-solvent interactions:

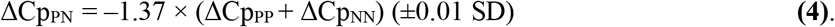

The overall consequence is that enthalpy stabilizes the folded state more at higher temperatures. The joint contribution of the two solvation terms (negative ΔCp_PN_ + positive ΔCp_NN_) to ΔCp_fol_ is negative, but the sign of the heat capacity change is specifically determined by protein-solvent interactions.

From the perspective of the forces acting on protein folding (**Figure 5b**), the contributions of van der Waals, coulombic and bonded interactions to ΔCp_fol_ are dissimilar. Van der Waals interactions show little dependence on temperature and, therefore, their contribution to ΔCp_fol_ is very small. Clearly, the major contribution to the negative value of ΔCp_fol_ comes from coulombic interactions, while bonded interactions also make a significant contribution. Thus, increasing temperature, markedly strengthens the stabilizing coulombic interactions and reduces the conformational strain of the folded conformation, which combines to produce a higher enthalpic stabilization of the native state. The sign of the folding heat capacity change is determined by coulombic interactions.

## Conclusions

Atomistic MD simulations in explicit solvent of native globular proteins and their unfolded ensembles reveal a coherent pattern of enthalpy and heat capacity changes when a polypeptide folds. The native conformation is enthalpically stabilized by similarly intense contributions from internal protein-protein and solvent-solvent interactions and destabilized by protein-solvent interactions. Van de Waals interactions mainly, but also coulombic interactions, stabilize the native conformation at the expense of introducing some conformational strain. The sign of the heat capacity change is set by the temperature dependence of protein-solvent interactions or, from the perspective of elementary contributions, by the temperature dependence of coulombic interactions. Changes in folding heat capacity but not in folding enthalpy appear to correlate well with protein length and with changes in protein solvent exposure upon folding.

## ASSOCIATED CONTENT

### Supplemental Information

#### SI TABLES

**SI Table 1**. Main elements of the MD setup, thermodynamic and solvating conditions used in the simulations of the proteins analyzed.

**SI Table 2**. Energy contributions to ΔH_fol_ and ΔCp_fol_ in simulations of barnase run with constraints on bonds involving Hydrogen atoms.

**SI Tables 3-6**. Data of energy terms extracted from MD simulations of barnase, SNase, apoFld and CI2, respectively, run with Charmm22-CMAP force field.

**SI Table 7**. Data of energy terms extracted from MD simulations of barnase run with Amber99SB-ILDN force field.

**SI Table 8**. Data of protein features (charges, isoelectric point, length, ΔSASA change upon folding and its polar and apolar components) and experimental values of ΔH_fol_ and ΔCp_fol_.

#### SI FIGURES

**SI Figure 1**. Linear correlation between calculated and experimentally determined ΔH_fol_ and ΔCp_fol_.

**SI Figure 2**. Relative contributions of molecular interactions to ΔH_fol_ and ΔCp_fol_.

**SI Figure 3**. Correlation between ΔSASA_apol_ and ΔSASA_pol_ and molecular and elementary contributions to ΔCp_fol_.

#### SI REFERENCES

### Author Information

ORCID

Juan J. Galano-Frutos: 0000-0002-1896-7805

Javier Sancho: 0000-0002-2879-9200

### Corresponding Author

Javier Sancho, Tel: +34 976 761 286.

### Author Contributions

J.S. conceived and directed the investigation. J.J.G-F. carried out and analysed the Molecular Dynamics simulations. J.J.G-F. and J.S. analysed data and wrote the manuscript.

### Funding Sources

This work was supported by grants PID2019-107293GB-I00 and PDC2021-121341-I00 (MICINN, Spain) and E45_20R (Gobierno de Aragón, Spain).

### Notes

The authors declare no conflict of interest.

## Acknowledgements

We thank the Biocomputation and Complex Systems Physics Institute (BIFI) of the University of Zaragoza and the Red Española de Supercomputación (RES) for computing facilities granted to perform Molecular Dynamics simulations.

## Abbreviations

IS: <u>I</u>onic <u>S</u>trength
LN: <u>L</u>ennard-<u>J</u>ones
MD: <u>M</u>olecular <u>D</u>ynamics
NMR: <u>N</u>uclear <u>M</u>agnetic <u>R</u>esonance
NN: <u>N</u>on-protein-<u>N</u>on-protein
PP: <u>P</u>rotein-<u>P</u>rotein
PN: <u>P</u>rotein-<u>N</u>on-protein
R_g_: Radius of gyration
SASA: <u>S</u>olvent-Accessible <u>S</u>urface <u>A</u>rea
SAXS: <u>S</u>mall-<u>A</u>ngle <u>X</u>-ray <u>S</u>cattering

## References

1. Anfinsen, C. B. Principles that govern the folding of protein chains. Science. 181, 223–230 (1973).

2. Dill, K. A. & MacCallum, J. L. The protein-folding problem, 50 years on. Science. 338, 1042–1046 (2012).

3. Huang, P. S., Boyken, S. E. & Baker, D. The coming of age of de novo protein design. Nature 537, 320–327 (2016).

4. Stein, A., Fowler, D. M., Hartmann-Petersen, R. & Lindorff-Larsen, K. Biophysical and Mechanistic models for disease-causing protein variants. Trends Biochem. Sci. 44, 575 (2019).

5. Sancho, J. The stability of 2-state, 3-state and more-state proteins from simple spectroscopic techniques… plus the structure of the equilibrium intermediates at the same time. Arch. Biochem. Biophys. 531, 4–13 (2013).

6. Lazaridis, T. & Karplus, M. Thermodynamics of protein folding: a microscopic view. Biophys. Chem. 100, 367–395 (2002).

7. Prabhu, N. V. & Sharp, K. A. Heat capacity in proteins. Annu. Rev. Phys. Chem. 56, 521–548 (2005).

8. Horovitz, A. Double-mutant cycles: a powerful tool for analyzing protein structure and function. Fold. Des. 1, R121–R126 (1996).

9. Gómez, J., Hilser, V. J., Xie, D. & Freire, E. The heat capacity of proteins. Proteins Struct. Funct. Bioinforma. 22, 404–412 (1995).

10. Campos, L. A., Cuesta-López, S., López-Llano, J., Falo, F. & Sancho, J. A Double-Deletion Method to Quantifying Incremental Binding Energies in Proteins from Experiment: Example of a Destabilizing Hydrogen Bonding Pair. Biophys. J. 88, 1311 (2005).

11. Bottaro, S. & Lindorff-Larsen, K. Biophysical experiments and biomolecular simulations: A perfect match? Science (80-.). 361, 355–360 (2018).

12. Piana, S., Lindorff-Larsen, K. & Shaw, D. E. Protein folding kinetics and thermodynamics from atomistic simulation. Proc. Natl. Acad. Sci. U. S. A. 109, 17845–17850 (2012).

13. Best, R. B. Atomistic molecular simulations of protein folding. Curr. Opin. Struct. Biol. 22, 52–61 (2012).

14. Piana, S., Robustelli, P., Tan, D., Chen, S. & Shaw, D. E. Development of a Force Field for the Simulation of Single-Chain Proteins and Protein-Protein Complexes. J. Chem. Theory Comput. 16, 2494–2507 (2020).

15. Robustelli, P., Piana, S. & Shaw, D. E. Developing a molecular dynamics force field for both folded and disordered protein states. Proc. Natl. Acad. Sci. U. S. A. 115, E4758–E4766 (2018).

16. Kamenik, A. S. et al. Polarizable and non-polarizable force fields: Protein folding, unfolding, and misfolding. J. Chem. Phys. 153, 185102 (2020).

17. Cui, X., Liu, H., Rehman, A. U. & Chen, H. F. Extensive evaluation of environment-specific force field for ordered and disordered proteins. Phys. Chem. Chem. Phys. 23, 12127–12136 (2021).

18. Prabhu, N. V. & Sharp, K. A. Heat capacity in proteins. Annu. Rev. Phys. Chem. 56, 521–548 (2005).

19. Robertson, A. D. & Murphy, K. P. Protein structure and the energetics of protein stability. Chem. Rev. 97, 1251–1267 (1997).

20. Newberry, R. W. & Raines, R. T. Secondary Forces in Protein Folding. ACS Chem. Biol. 14, 1677 (2019).

21. Galano-Frutos, J. J. & Sancho, J. Accurate Calculation of Barnase and SNase Folding Energetics Using Short Molecular Dynamics Simulations and an Atomistic Model of the Unfolded Ensemble: Evaluation of Force Fields and Water Models. J. Chem. Inf. Model. 59, 4350–4360 (2019).

22. Galano-Frutos, J. J., Sancho, J. & Nerín-Fonz, F. Calculation of Protein Folding Thermodynamics using Molecular Dynamics Simulations. bioRxiv Preprint (2023) doi:https://doi.org/10.1101/2023.01.21.525008.

23. Estrada, J., Bernadó, P., Blackledge, M. & Sancho, J. ProtSA: A web application for calculating sequence specific protein solvent accessibilities in the unfolded ensemble. BMC Bioinformatics 10, 1–8 (2009).

24. Piana, S., Klepeis, J. L. & Shaw, D. E. Assessing the accuracy of physical models used in protein-folding simulations: quantitative evidence from long molecular dynamics simulations. Curr. Opin. Struct. Biol. 24, 98–105 (2014).

25. Zerze, G. H., Zheng, W., Best, R. B. & Mittal, J. Evolution of All-Atom Protein Force Fields to Improve Local and Global Properties. J. Phys. Chem. Lett. 10, 2227–2234 (2019).

26. Best, R. B., Zheng, W. & Mittal, J. Balanced protein-water interactions improve properties of disordered proteins and non-specific protein association. J. Chem. Theory Comput. 10, 5113–5124 (2014).

27. Piana, S., Donchev, A. G., Robustelli, P. & Shaw, D. E. Water dispersion interactions strongly influence simulated structural properties of disordered protein states. J. Phys. Chem. B 119, 5113–5123 (2015).

28. Hartley, R. W. & Barker, E. A. Amino-acid sequence of extracellular ribonuclease (barnase) of Bacillus amyloliquefaciens. Nat. New Biol. 235, 15–16 (1972).

29. Kullik, I., Giachino, P. & Fuchs, T. Deletion of the alternative sigma factor σ(B) in Staphylococcus aureus reveals its function as a global regulator of virulence genes. J. Bacteriol. 180, 4814–4820 (1998).

30. Fillat, M. F., Edmondson, D. E. & Gomez-Moreno, C. Structural and chemical properties of a flavodoxin from Anabaena PCC 7119. Biochim. Biophys. Acta - Protein Struct. Mol. Enzymol. 1040, 301–307 (1990).

31. Svendsen, I., Martin, B. & Jonassen, I. Characteristics of Hiproly barley III. Amino acid sequences of two lysine-rich proteins. Carlsberg Res. Commun. 45, 79–85 (1980).

32. Martin, C., Richard, V., Salem, M., Hartley, R. & Mauguen, Y. Refinement and structural analysis of barnase at 1.5 A resolution. Acta Crystallogr. D. Biol. Crystallogr. 55, 386–398 (1999).

33. Cotton, F. A., Hazen, E. E. & Legg, M. J. Staphylococcal nuclease: Proposed mechanism of action based on structure of enzyme—thymidine 3′,5′-bisphosphate—calcium ion complex at 1.5-Å resolution. Proc. Natl. Acad. Sci. U. S. A. 76, 2551 (1979).

34. Genzor, C. G., Perales-Alcon, A., Sancho, J. & Romero, A. Closure of a tyrosine/tryptophan aromatic gate leads to a compact fold in apo flavodoxin. Nat. Struct. Biol. 3, 329–332 (1996).

35. McPhalen, C. A. & James, M. N. Crystal and Molecular Structure of the Serine Proteinase Inhibitor CI-2 from Barley Seeds. Biochemistry 26, 261–269 (1987).

36. Jorgensen, W. L., Chandrasekhar, J., Madura, J. D., Impey, R. W. & Klein, M. L. Comparison of simple potential functions for simulating liquid water. J. Chem. Phys. 79, 926–935 (1983).

37. Mackerell, A. D., Feig, M. & Brooks, C. L. Extending the treatment of backbone energetics in protein force fields: Limitations of gas-phase quantum mechanics in reproducing protein conformational distributions in molecular dynamics simulation. J. Comput. Chem. 25, 1400–1415 (2004).

38. Van Der Spoel, D. et al. GROMACS: Fast, flexible, and free. Journal of Computational Chemistry vol. 26 1701–1718 (2005).

39. Becktel, W. J. & Schellman, J. A. Protein stability curves. Biopolymers 26, 1859–1877 (1987).

40. Lindorff-Larsen, K. et al. Improved side-chain torsion potentials for the Amber ff99SB protein force field. Proteins Struct. Funct. Bioinforma. 78, 1950–1958 (2010).

41. Shao, Q., Yang, L. & Zhu, W. Selective enhanced sampling in dihedral energy facilitates overcoming the dihedral energy increase in protein folding and accelerates the searching for protein native structure. Phys. Chem. Chem. Phys. 21, 10423–10435 (2019).

42. Baldwin, R. L. Temperature dependence of the hydrophobic interaction in protein folding. Proc. Natl. Acad. Sci. U. S. A. 83, 8069 (1986).

43. Privalov, P. L. & Gill, S. J. Stability of Protein Structure and Hydrophobic Interaction. Adv. Protein Chem. 39, 191–234 (1988).

44. Murphy, K. P., Privalov, P. L. & Gill, S. J. Common Features of Protein Unfolding and Dissolution of Hydrophobic Compounds. Science (80-.). 247, 559–561 (1990).

45. Makhatadze, G. I. & Privalov, P. L. Heat capacity of proteins: I. Partial molar heat capacity of individual amino acid residues in aqueous solution: Hydration effect. J. Mol. Biol. 213, 375–384 (1990).

46. Murphy, K. P. & Gill, S. J. Solid model compounds and the thermodynamics of protein unfolding. J. Mol. Biol. 222, 699–709 (1991).

47. Hilser, V. J., Gómez, J. & Freire, E. The enthalpy change in protein folding and binding: refinement of parameters for structure-based calculations. Proteins 26, 123–133 (1996).

48. Privalov, P. L. & Makhatadze, G. I. Contribution of hydration and non-covalent interactions to the heat capacity effect on protein unfolding. J. Mol. Biol. 224, 715–723 (1992).

49. Madan, B. & Sharp, K. Heat Capacity Changes Accompanying Hydrophobic and Ionic Solvation: A Monte Carlo and Random Network Model Study. J. Phys. Chem. 100, 7713–7721 (1996).

50. Ravikumar, Y., Nadarajan, S. P., Hyeon Yoo, T., Lee, C. S. & Yun, H. Unnatural amino acid mutagenesis-based enzyme engineering. Trends Biotechnol. 33, 462–470 (2015).

51. Chin, J. W. Expanding and reprogramming the genetic code. Nat. 2017 5507674 550, 53–60 (2017).

52. Xavier, J. S. et al. ThermoMutDB: a thermodynamic database for missense mutations. Nucleic Acids Res. 49, D475 (2021).

53. Nikam, R., Kulandaisamy, A., Harini, K., Sharma, D. & Michael Gromiha, M. ProThermDB: thermodynamic database for proteins and mutants revisited after 15 years. Nucleic Acids Res. 49, D420 (2021).

54. García-Cebollada, H., López, A. & Sancho, J. Protposer: The web server that readily proposes protein stabilizing mutations with high PPV. Comput. Struct. Biotechnol. J. 20, 2415 (2022).

55. Dehouck, Y., Kwasigroch, J. M., Gilis, D. & Rooman, M. PoPMuSiC 2.1: A web server for the estimation of protein stability changes upon mutation and sequence optimality. BMC Bioinformatics 12, 1–12 (2011).

56. Lei, H., Wu, C., Liu, H. & Duan, Y. Folding free-energy landscape of villin headpiece subdomain from molecular dynamics simulations. Proc. Natl. Acad. Sci. U. S. A. 104, 4925–4930 (2007).

57. Ferina, J. & Daggett, V. Visualizing Protein Folding and Unfolding. J. Mol. Biol. 431, 1540–1564 (2019).

58. Best, R. B. Analysis of Molecular Dynamics Simulations of Protein Folding. Methods Mol. Biol. 2376, 317–329 (2022).

59. Lindorff-Larsen, K., Piana, S., Dror, R. O. & Shaw, D. E. How fast-folding proteins fold. Science (80-.). 334, 517–520 (2011).

60. Mackerell, A. D. Empirical force fields for biological macromolecules: Overview and issues. J. Comput. Chem. 25, 1584–1604 (2004).

61. Baker, D. What has de novo protein design taught us about protein folding and biophysics? Protein Sci. 28, 678–683 (2019).

