## Supplemental Material for "Molecular Interactions and Forces that Make Proteins Stable: A Quantitative Inventory from Atomistic Molecular Dynamics Simulations"

### Content

#### SI TABLES

**SI Table 1.** Main elements of the MD setup, thermodynamic and solvating conditions used in the simulations of the proteins analyzed.

**SI Table 2.** Energy contributions to $\Delta H_{fol}$ and $\Delta Cp_{fol}$ in simulations of barnase run with constraints on bonds involving Hydrogen atoms.

**SI Tables 3-6.** Data of energy terms extracted from MD simulations of barnase, SNase, apoFld and CI2, respectively, run with Charmm22-CMAP force field.

**SI Table 7.** Data of energy terms extracted from MD simulations of barnase run with Amber99SB-ILDN force field.

**SI Table 8.** Data of protein features (charges, isoelectric point, length, ΔSASA change upon folding and its polar and apolar components) and experimental values of $\Delta H_{fol}$ and $\Delta Cp_{fol}$.

#### SI FIGURES

**SI Figure 1.** Linear correlation between calculated and experimentally determined $\Delta H_{fol}$ and $\Delta Cp_{fol}$.

**SI Figure 2.** Relative contributions of molecular interactions to $\Delta H_{fol}$ and $\Delta Cp_{fol}$.

**SI Figure 3.** Correlation between $\Delta SASA_{apol}$ and $\Delta SASA_{pol}$ and molecular and elementary contributions to $\Delta Cp_{fol}$.

#### SI REFERENCES

**SI Table 1.** Main elements of the MD setup, thermodynamic and solvating conditions used in the simulations of the proteins analyzed

| **Protein & Force field** | **MD setup[a] and solvating conditions** | | | | |
|---|---|---|---|---|---|
| | **Temperature and pressure** | **Number of replicas** | **Simulated pH & Ionic Strength (mM)** | **Number & type of ions** | **No. water molecules in the simulated box** |
| barnase (Charmm)[b] | T = 295,315,335 K<br>P = 1 atm | Folded (40),<br>Unfolded (100) | pH ~4.1,<br>IS = 4.0 | 4 $Na^+$,<br>11 $Cl^-$ | 63,892 |
| barnase (Amber)[c] | T = 295,315,335 K<br>P = 1 atm | Folded (10),<br>Unfolded (40) | pH ~4.1,<br>IS = 6.0 | 7 $Cl^-$ | 63,921 |
| SNase (Charmm)[b] | T = 307,317,327 K<br>P = 1 atm | Folded (40),<br>Unfolded (100) | pH 7.0,<br>IS = 5.0 | 10 $Cl^-$ | 115,602 |
| apoFld (Charmm)[b] | T = 305,320,335 K<br>P = 1 atm | Folded (40),<br>Unfolded (100) | pH 7.0,<br>IS = 17.0 | 51 $Na^+$,<br>34 $Cl^-$ | 116,732 |
| CI2 (Charmm)[b] | T = 320,335,350 K<br>P = 1 atm | Folded (40),<br>Unfolded (100) | pH 3.0,<br>IS = 1.9 | 1 $Na^+$,<br>6 $Cl^-$ | 30,950 |

[a] All simulations were performed in the NPT ensemble using the v-rescale thersmostat and the Parrinello-Rahman barostat.

[b] Simulations performed with Charmm22-CMAP[1] force field (Gromacs 2020 package[2]), reported in Galano-Frutos et. al.[3]. All proteins were simulated under constraints on all bonds but, for comparison purposes, barnase was also simulated under constraints on only bonds involving hydrogen atoms (see **SI Table 2**). Additional MD setup and solvating conditions details are given in the **Supplementary Material** of Galano-Frutos et. al.[3].

[c] Simulations performed with Amber99SB-ILDN[4] force field (Gromacs 4.6.7 package[2]) reported in Galano-Frutos et. al.[5] Additional MD setup and solvating conditions details are given in this article[5].

**SI Table 2.** Energy contributions to $\Delta H_{fol}$ and $\Delta Cp_{fol}$ in simulations of barnase run with constraints on bonds involving hydrogen atoms[a]

| **Contributor**[b] | **Folding Energy Change** | | | **ΔCp** | $R^2$ |
|---|---|---|---|---|---|
| | **Temperature** | | | | |
| | **295 K** | **315 K** | **335 K** | | |
| $\Delta E_{PP}$ | −3,044 | −2,950 | −2,769 | 6.87 | 0.97 |
| $\Delta E_{PN}$ | 5,164 | 4,866 | 4,413 | −18.76 | 0.99 |
| $\Delta E_{NN}$ | −2,457 | −2,344 | −2,158 | 7.47 | 0.98 |
| $\Delta E^{Coul}$ | −143 | −200 | −262 | −2.97 | 1.00 |
| $\Delta E^{LJ}$ | −263 | −276 | −273 | −0.25 | 0.57 |
| $\Delta E^{Bonded}$ | 68 | 48 | 21 | −1.19 | 0.99 |
| ΔE | −337 | −428 | −514 | −4.42 | 1.00 |
| $\Delta E^{kin}$ | 7 | 9 | 9 | 0.05 | 0.75 |
| Δ(pV) | 0 | 0 | 0 | NA | NA |
| ΔH | −330 | −419 | −505 | −4.36 | 1.00 |

[a] Data from simulations run with Charmm22-CMAP[1] force field, Tip3p[6] water model, and constraints imposed on bonds involving hydrogen atoms.

[b] Equivalences: $\Delta E_{PP} = \Delta E_{PP}^{LJ} + \Delta E_{PP}^{Coul} + \Delta E^{Bonded}$; $\Delta E_{PN} = \Delta E_{PN}^{LJ} + \Delta E_{PN}^{Coul}$; $\Delta E_{NN} = \Delta E_{NN}^{LJ} + \Delta E_{NN}^{Coul} + \Delta E_{NN}^{Coul-recip} + \Delta E_{NN}^{Disp-Corr}$.

**SI Table 3.** Data of energy terms (Charmm22-CMAP force field) extracted from the MD simulations of barnase performed at three temperatures

| Force field terms[a] | 295 K | | | | | | 315 K | | | | | | 335 K | | | | | |
|---|---|---|---|---|---|---|---|---|---|---|---|---|---|---|---|---|---|---|
| | FOLD[b] | | UNF[c] | | FOLD-UNF[d] | | UNF[c] | | FOLD[b] | | FOLD-UNF[d] | | FOLD[b] | | UNF[c] | | FOLD-UNF[d] | |
| | Ave | SD | Ave | SD | Ave | SD | Ave | SD | Ave | SD | Ave | SD | Ave | SD | Ave | SD | Ave | SD |
| U-B | 3,716 | 6 | 3,690 | 30 | 25 | 36 | 3,889 | 10 | 3,865 | 26 | 24 | 37 | 4,069 | 8 | 4,046 | 26 | 23 | 34 |
| Proper-Dih | 2,466 | 10 | 2,423 | 46 | 43 | 56 | 2,514 | 11 | 2,490 | 44 | 23 | 55 | 2,559 | 12 | 2,556 | 42 | 3 | 54 |
| Improp-Dih | 227 | 1 | 225 | 1 | 2 | 2 | 240 | 1 | 239 | 1 | 1 | 2 | 254 | 1 | 254 | 1 | 0 | 2 |
| CMAP-Dih | −693 | 6 | −711 | 52 | 18 | 58 | −692 | 10 | −705 | 51 | 13 | 61 | −682 | 7 | −705 | 51 | 24 | 59 |
| LJ-14 | 1,778 | 6 | 1,739 | 13 | 40 | 18 | 1,790 | 4 | 1,749 | 11 | 41 | 15 | 1,800 | 4 | 1,759 | 11 | 42 | 15 |
| Coul-14 | 17,219 | 58 | 17,181 | 129 | 38 | 187 | 17,211 | 53 | 17,172 | 118 | 39 | 172 | 17,180 | 70 | 17,147 | 111 | 32 | 181 |
| LJ-SR | 396,558 | 21 | 396,876 | 37 | −318 | 58 | 375,347 | 25 | 375,665 | 36 | −318 | 61 | 355,549 | 21 | 355,875 | 37 | −326 | 58 |
| Disp-corr | −11,293 | 0 | −11,293 | 0 | 0 | 1 | −11,088 | 0 | −11,087 | 0 | −1 | 1 | −10,862 | 0 | −10,861 | 0 | −1 | 1 |
| Coul-SR | −2,739,844 | 50 | −2,739,747 | 77 | −97 | 127 | −2,650,641 | 40 | −2,650,472 | 75 | −168 | 114 | −2,563,747 | 55 | −2,563,539 | 63 | −208 | 118 |
| Coul-recip | −263,633 | 25 | −263,554 | 57 | −79 | 82 | −262,124 | 20 | −262,037 | 52 | −87 | 73 | −260,499 | 30 | −260,407 | 50 | −92 | 80 |
| Potential | −2,593,503 | 29 | −2,593,172 | 46 | −331 | 75 | −2,523,554 | 36 | −2,523,121 | 47 | −432 | 83 | −2,454,380 | 43 | −2,453,876 | 43 | −503 | 85 |
| Kin-Energy | 474,354 | 6 | 474,358 | 7 | −3 | 13 | 506,513 | 9 | 506,519 | 7 | −6 | 16 | 538,675 | 8 | 538,680 | 6 | −5 | 14 |
| Total-Energy | −2,119,148 | 28 | −2,118,815 | 45 | −333 | 73 | −2,017,040 | 33 | −2,016,603 | 47 | −437 | 80 | −1,915,706 | 43 | −1,915,196 | 43 | −509 | 86 |
| Temp | 295 | 0 | 295 | 0 | 0 | 0 | 315 | 0 | 315 | 0 | 0 | 0 | 335 | 0 | 335 | 0 | 0 | 0 |
| Press | 1 | 0 | 1 | 0 | 0 | 1 | 1 | 0 | 1 | 0 | 0 | 1 | 1 | 0 | 1 | 0 | 0 | 1 |
| Vol | 1,947 | 0 | 1,947 | 0 | 0 | 0 | 1,983 | 0 | 1,983 | 0 | 0 | 0 | 2,024 | 0 | 2,025 | 0 | 0 | 0 |
| Density | 992 | 0 | 992 | 0 | 0 | 0 | 974 | 0 | 974 | 0 | 0 | 0 | 955 | 0 | 954 | 0 | 0 | 0 |
| pV | 117 | 0 | 117 | 0 | 0 | 0 | 119 | 0 | 119 | 0 | 0 | 0 | 122 | 0 | 122 | 0 | 0 | 0 |
| Enthalpy | −2,119,031 | 29 | −2,118,697 | 45 | −334 | 74 | −2,016,921 | 33 | −2,016,483 | 48 | −438 | 81 | −1,915,583 | 43 | −1,915,074 | 42 | −509 | 85 |
| Coul-SR-P-P | −6,066 | 70 | −3,952 | 188 | −2,114 | 258 | −6,045 | 70 | −4,056 | 208 | −1,988 | 278 | −6,004 | 78 | −4,126 | 198 | −1,879 | 275 |
| LJ-SR-P-P | −2,854 | 15 | −1,693 | 99 | −1,161 | 114 | −2,836 | 12 | −1,705 | 108 | −1,131 | 119 | −2,809 | 12 | −1,714 | 100 | −1,095 | 111 |
| Coul-14-P-P | 17,219 | 58 | 17,181 | 129 | 38 | 187 | 17,211 | 53 | 17,172 | 118 | 39 | 172 | 17,180 | 70 | 17,147 | 111 | 32 | 181 |

**SI Table 3.** Continuation...

| Force field terms[a] | 295 K | | | | | | 315 K | | | | | | 335 K | | | | | |
|---|---|---|---|---|---|---|---|---|---|---|---|---|---|---|---|---|---|---|
| | FOLD[b] | | UNF[c] | | FOLD-UNF[d] | | UNF[c] | | FOLD[b] | | FOLD-UNF[d] | | FOLD[b] | | UNF[c] | | FOLD-UNF[d] | |
| | Ave | SD | Ave | SD | Ave | SD | Ave | SD | Ave | SD | Ave | SD | Ave | SD | Ave | SD | Ave | SD |
| LJ-14-P-P | 1,778 | 6 | 1,739 | 13 | 40 | 18 | 1,790 | 4 | 1,749 | 11 | 41 | 15 | 1,800 | 4 | 1,759 | 11 | 42 | 15 |
| Coul-SR-P-N | −8,472 | 138 | −12,818 | 343 | 4,346 | 481 | −8,260 | 125 | −12,299 | 399 | 4,038 | 524 | −8,070 | 154 | −11,832 | 375 | 3,762 | 529 |
| LJ-SR-P-N | −683 | 19 | −1,598 | 92 | 915 | 112 | −665 | 13 | −1,534 | 96 | 869 | 110 | −645 | 15 | −1,467 | 88 | 822 | 103 |
| Coul-SR-N-N | −2,725,307 | 80 | −2,722,977 | 170 | −2,330 | 249 | −2,636,336 | 71 | −2,634,118 | 189 | −2,218 | 260 | −2,549,673 | 84 | −2,547,581 | 180 | −2,092 | 264 |
| LJ-SR-N-N | 400,095 | 21 | 400,167 | 28 | −72 | 49 | 378,849 | 16 | 378,905 | 24 | −56 | 40 | 359,003 | 18 | 359,056 | 25 | −53 | 43 |

[a] Energy terms issued by Gromacs (from the .edr file) in simulations of barnase with Charmm22-CMAP[1] force field and Tip3p[6] water model.

[b] Averages (Ave) and standard deviations (SD) calculated from the forty (40) replicas simulated for the protein folded (FOLD) state (reported in Galano-Frutos et. al.)[3].

[c] Averages (Ave) and standard deviations (SD) calculated from the hundred (100) unfolded structures simulated for the protein unfolded (UNF) state (reported in Galano-Frutos et. al.)[3].

**SI Table 4.** Data of energy terms (Charmm22-CMAP force field) extracted from the MD simulations of SNase performed at three temperatures

| **Force field terms[a]** | **307 K** | | | | | | **317 K** | | | | | | **327 K** | | | | | |
|---|---|---|---|---|---|---|---|---|---|---|---|---|---|---|---|---|---|---|
| | **FOLD[b]** | | **UNF[c]** | | **FOLD-UNF[d]** | | **UNF[c]** | | **FOLD[b]** | | **FOLD-UNF[d]** | | **FOLD[b]** | | **UNF[c]** | | **FOLD-UNF[d]** | |
| | **Ave** | **SD** | **Ave** | **SD** | **Ave** | **SD** | **Ave** | **SD** | **Ave** | **SD** | **Ave** | **SD** | **Ave** | **SD** | **Ave** | **SD** | **Ave** | **SD** |
| U-B | 5,550 | 14 | 5,469 | 32 | 81 | 46 | 5,677 | 18 | 5,598 | 33 | 79 | 51 | 5,800 | 19 | 5,728 | 31 | 72 | 50 |
| Proper-Dih | 3,402 | 23 | 3,332 | 55 | 70 | 77 | 3,439 | 19 | 3,374 | 54 | 64 | 74 | 3,467 | 20 | 3,417 | 51 | 50 | 72 |
| Improper-Dih | 323 | 1 | 322 | 1 | 2 | 3 | 333 | 1 | 332 | 1 | 1 | 2 | 342 | 1 | 342 | 2 | 0 | 3 |
| CMAP-Dih | −906 | 22 | −920 | 64 | 14 | 86 | −918 | 16 | −914 | 64 | −3 | 81 | −920 | 18 | −910 | 59 | −11 | 77 |
| LJ-14 | 1,924 | 9 | 1,882 | 14 | 42 | 23 | 1,934 | 7 | 1,887 | 15 | 47 | 22 | 1,945 | 9 | 1,895 | 14 | 50 | 24 |
| Coul-14 | 30,679 | 91 | 30,983 | 131 | −304 | 223 | 30,697 | 81 | 30,978 | 127 | −282 | 208 | 30,713 | 86 | 30,983 | 129 | −270 | 215 |
| LJ-SR | 695,504 | 35 | 695,949 | 49 | −445 | 84 | 676,790 | 32 | 677,241 | 45 | −451 | 77 | 658,693 | 28 | 659,142 | 46 | −449 | 74 |
| Disp-corr | −20,094 | 1 | −20,095 | 1 | 1 | 1 | −19,902 | 1 | −19,903 | 1 | 1 | 1 | −19,702 | 0 | −19,702 | 1 | 0 | 1 |
| Coul-SR | −4,853,618 | 85 | −4,853,938 | 96 | 320 | 181 | −4,773,784 | 81 | −4,774,040 | 84 | 256 | 164 | −4,694,948 | 78 | −4,695,173 | 86 | 225 | 164 |
| Coul-recip | −466,063 | 43 | −466,113 | 55 | 51 | 98 | −464,681 | 35 | −464,713 | 54 | 32 | 89 | −463,246 | 38 | −463,272 | 54 | 26 | 91 |
| Potential | −4,603,300 | 55 | −4,603,128 | 69 | −171 | 124 | −4,540,415 | 70 | −4,540,158 | 55 | −257 | 126 | −4,477,855 | 60 | −4,477,550 | 52 | −305 | 112 |
| Kin-Energy | 891,376 | 9 | 891,374 | 10 | 2 | 19 | 920,407 | 9 | 920,408 | 9 | −1 | 18 | 949,441 | 9 | 949,445 | 9 | −4 | 19 |
| Total-Energy | −3,711,923 | 56 | −3,711,754 | 69 | −168 | 125 | −3,620,009 | 67 | −3,619,750 | 54 | −258 | 121 | −3,528,414 | 58 | −3,528,104 | 53 | −309 | 111 |
| Temp | 307 | 0 | 307 | 0 | 0 | 0 | 317 | 0 | 317 | 0 | 0 | 0 | 327 | 0 | 327 | 0 | 0 | 0 |
| Press | 1 | 0 | 1 | 0 | 0 | 0 | 1 | 0 | 1 | 0 | 0 | 0 | 1 | 0 | 1 | 0 | 0 | 0 |
| Vol | 3,555 | 0 | 3,555 | 0 | 0 | 0 | 3,589 | 0 | 3,589 | 0 | 0 | 0 | 3,626 | 0 | 3,626 | 0 | 0 | 0 |
| Density | 981 | 0 | 981 | 0 | 0 | 0 | 971 | 0 | 972 | 0 | 0 | 0 | 962 | 0 | 962 | 0 | 0 | 0 |
| pV | 214 | 0 | 214 | 0 | 0 | 0 | 216 | 0 | 216 | 0 | 0 | 0 | 218 | 0 | 218 | 0 | 0 | 0 |
| Enthalpy | −3,711,709 | 56 | −3,711,540 | 69 | −169 | 125 | −3,619,793 | 66 | −3,619,535 | 54 | −258 | 121 | −3,528,195 | 59 | −3,527,886 | 52 | −309 | 112 |
| Coul-SR-P-P | −8,893 | 146 | −6,206 | 258 | −2,687 | 404 | −8,921 | 172 | −6,253 | 262 | −2,667 | 434 | −8,911 | 155 | −6,377 | 281 | −2,534 | 435 |
| LJ-SR-P-P | −3,765 | 33 | −2,225 | 117 | −1,540 | 150 | −3,749 | 35 | −2,224 | 118 | −1,525 | 153 | −3,732 | 45 | −2,246 | 120 | −1,486 | 165 |
| Coul-14-P-P | 30,679 | 91 | 30,983 | 131 | −304 | 223 | 30,697 | 81 | 30,978 | 127 | −282 | 208 | 30,713 | 86 | 30,983 | 129 | −270 | 215 |

**SI Table 4.** Continuation...

| | 295 K | | | | | | 315 K | | | | | | 335 K | | | | | |
|---|---|---|---|---|---|---|---|---|---|---|---|---|---|---|---|---|---|---|
| **Force field terms[a]** | **FOLD[b]** | | **UNF[c]** | | **FOLD-UNF[d]** | | **UNF[c]** | | **FOLD[b]** | | **FOLD-UNF[d]** | | **FOLD[b]** | | **UNF[c]** | | **FOLD-UNF[d]** | |
| | **Ave** | **SD** | **Ave** | **SD** | **Ave** | **SD** | **Ave** | **SD** | **Ave** | **SD** | **Ave** | **SD** | **Ave** | **SD** | **Ave** | **SD** | **Ave** | **SD** |
| LJ-14-P-P | 1,924 | 9 | 1,882 | 14 | 42 | 23 | 1,934 | 7 | 1,887 | 15 | 47 | 22 | 1,945 | 9 | 1,895 | 14 | 50 | 24 |
| Coul-SR-P-N | −13,557 | 267 | −19,780 | 515 | 6,224 | 782 | −13,345 | 342 | −19,445 | 526 | 6,101 | 868 | −13,213 | 326 | −18,982 | 561 | 5,769 | 887 |
| LJ-SR-P-N | −755 | 35 | −1,957 | 110 | 1,202 | 145 | −759 | 39 | −1,918 | 103 | 1,159 | 142 | −738 | 37 | −1,865 | 108 | 1,127 | 145 |
| Coul-SR-N-N | −4,831,168 | 150 | −4,827,951 | 235 | −3,217 | 385 | −4,751,519 | 153 | −4,748,341 | 252 | −3,177 | 405 | −4,672,822 | 151 | −4,669,814 | 255 | −3,007 | 405 |
| LJ-SR-N-N | 700,024 | 29 | 700,132 | 41 | −108 | 70 | 681,298 | 23 | 681,383 | 31 | −85 | 54 | 663,163 | 26 | 663,254 | 33 | −91 | 59 |

[a] Energy terms issued by GROMACS (from the .edr file) in simulations of nuclease with Charmm22-CMAP[1] force field and Tip3p[6] water model.

[b] Averages (Ave) and standard deviations (SD) calculated from the forty (40) replicas simulated for the protein folded (FOLD) state (reported in Galano-Frutos et. al.)[3].

[c] Averages (Ave) and standard deviations (SD) calculated from the hundred (100) unfolded structures simulated for the protein unfolded (UNF) state (reported in Galano-Frutos et. al.)[3].

**SI Table 5.** Data of energy terms (Charmm22-CMAP force field) extracted from the MD simulations of apoFld performed at three temperatures

| Force field terms[a] | 305 K | | | | | | 320 K | | | | | | 335 K | | | | | |
|---|---|---|---|---|---|---|---|---|---|---|---|---|---|---|---|---|---|---|
| | FOLD[b] | | UNF[c] | | FOLD-UNF[d] | | UNF[c] | | FOLD[b] | | FOLD-UNF[d] | | FOLD[b] | | UNF[c] | | FOLD-UNF[d] | |
| | Ave | SD | Ave | SD | Ave | SD | Ave | SD | Ave | SD | Ave | SD | Ave | SD | Ave | SD | Ave | SD |
| U-B | 5,705 | 11 | 5,685 | 34 | 20 | 45 | 5,906 | 17 | 5,885 | 35 | 21 | 53 | 6,101 | 13 | 6,090 | 35 | 11 | 48 |
| Proper-Dih | 3,518 | 13 | 3,520 | 52 | −2 | 65 | 3,559 | 17 | 3,589 | 55 | −31 | 72 | 3,602 | 16 | 3,660 | 51 | −59 | 67 |
| Improper-Dih | 389 | 1 | 383 | 2 | 7 | 3 | 406 | 1 | 401 | 2 | 5 | 3 | 422 | 1 | 419 | 2 | 3 | 3 |
| CMAP-Dih | −982 | 15 | −1,141 | 58 | 159 | 72 | −982 | 16 | −1,131 | 66 | 149 | 82 | −971 | 20 | −1,134 | 62 | 163 | 82 |
| LJ-14 | 2,669 | 6 | 2,574 | 15 | 96 | 22 | 2,684 | 6 | 2,586 | 17 | 99 | 23 | 2,700 | 7 | 2,599 | 16 | 101 | 23 |
| Coul-14 | 36,443 | 85 | 36,624 | 140 | −181 | 225 | 36,435 | 82 | 36,627 | 156 | −192 | 238 | 36,476 | 78 | 36,628 | 149 | −152 | 227 |
| LJ-SR | 827,642 | 51 | 828,117 | 50 | −475 | 101 | 794,491 | 48 | 794,975 | 48 | −484 | 96 | 762,979 | 43 | 763,444 | 58 | −465 | 101 |
| Disp-corr | −23,776 | 2 | −23,777 | 1 | 1 | 2 | −23,435 | 2 | −23,436 | 1 | 1 | 2 | −23,070 | 2 | −23,070 | 1 | 0 | 2 |
| Coul-SR | −5,761,951 | 129 | −5,762,184 | 96 | 234 | 226 | −5,620,404 | 141 | −5,620,553 | 103 | 149 | 244 | −5,481,545 | 127 | −5,481,519 | 89 | −26 | 216 |
| Coul-recip | −552,621 | 37 | −552,576 | 63 | −44 | 100 | −550,146 | 37 | −550,098 | 69 | −48 | 106 | −547,558 | 35 | −547,484 | 66 | −74 | 101 |
| Potential | −546,2960 | 94 | −5,462,776 | 63 | −184 | 157 | −5,351,485 | 100 | −5,351,155 | 64 | −329 | 163 | −5,240,865 | 99 | −5,240,368 | 64 | −497 | 163 |
| Kin-Energy | 1,046,774 | 9 | 1,046,775 | 8 | −1 | 17 | 1,098,256 | 6 | 1,098,257 | 9 | 0 | 16 | 1,149,738 | 9 | 1,149,738 | 8 | 0 | 17 |
| Total-Energy | −4,416,187 | 94 | −4,416,001 | 64 | −186 | 158 | −4,253,229 | 99 | −4,252,899 | 63 | −330 | 162 | −4,091,126 | 96 | −4,090,629 | 64 | −497 | 161 |
| Temp | 305 | 0 | 305 | 0 | 0 | 0 | 320 | 0 | 320 | 0 | 0 | 0 | 335 | 0 | 335 | 0 | 0 | 0 |
| Press | 1 | 1 | 1 | 0 | −1 | 1 | 1 | 1 | 1 | 1 | 0 | 2 | 1 | 1 | 1 | 0 | 0 | 1 |
| Vol | 4,194 | 0 | 4,194 | 0 | 0 | 0 | 4,255 | 0 | 4,255 | 0 | 0 | 0 | 4,323 | 0 | 4,323 | 0 | 0 | 0 |
| Density | 983 | 0 | 983 | 0 | 0 | 0 | 969 | 0 | 969 | 0 | 0 | 0 | 954 | 0 | 954 | 0 | 0 | 0 |
| pV | 253 | 0 | 253 | 0 | 0 | 0 | 256 | 0 | 256 | 0 | 0 | 0 | 260 | 0 | 260 | 0 | 0 | 0 |
| Enthalpy | −4,415,934 | 93 | −4,415,748 | 63 | −186 | 156 | −4,252,972 | 98 | −4,252,642 | 62 | −330 | 160 | −4,090,866 | 96 | −4,090,369 | 64 | −497 | 161 |
| Coul-SR-P-P | −10,575 | 95 | −7,818 | 313 | −2,758 | 408 | −10,483 | 133 | −7,981 | 333 | −2,502 | 466 | −10,489 | 118 | −8,134 | 337 | −2,355 | 455 |
| LJ-SR-P-P | −4,627 | 34 | −2,713 | 138 | −1,913 | 172 | −4,592 | 37 | −2,718 | 158 | −1,874 | 196 | −4,549 | 32 | −2,730 | 160 | −1,820 | 193 |
| Coul-14-P-P | 36,443 | 85 | 36,624 | 140 | −181 | 225 | 36,435 | 82 | 36,627 | 156 | −192 | 238 | 36,476 | 78 | 36,628 | 149 | −152 | 227 |

**SI Table 5.** Continuation...

| Force field terms[a] | 295 K | | | | | | 315 K | | | | | | 335 K | | | | | |
|---|---|---|---|---|---|---|---|---|---|---|---|---|---|---|---|---|---|---|
| | FOLD[b] | | UNF[c] | | FOLD-UNF[d] | | UNF[c] | | FOLD[b] | | FOLD-UNF[d] | | FOLD[b] | | UNF[c] | | FOLD-UNF[d] | |
| | **Ave** | **SD** | **Ave** | **SD** | **Ave** | **SD** | **Ave** | **SD** | **Ave** | **SD** | **Ave** | **SD** | **Ave** | **SD** | **Ave** | **SD** | **Ave** | **SD** |
| LJ-14-P-P | 2,669 | 6 | 2,574 | 15 | 96 | 22 | 2,684 | 6 | 2,586 | 17 | 99 | 23 | 2,700 | 7 | 2,599 | 16 | 101 | 23 |
| Coul-SR-P-N | −16,044 | 180 | −22,568 | 594 | 6,524 | 773 | −15,969 | 245 | −21,853 | 647 | 5,885 | 892 | −15,706 | 236 | −21,151 | 673 | 5,444 | 909 |
| LJ-SR-P-N | −282 | 27 | −1,876 | 129 | 1,593 | 156 | −271 | 27 | −1,825 | 143 | 1,554 | 170 | −274 | 26 | −1,762 | 141 | 1,489 | 167 |
| Coul-SR-N-N | −5,735,333 | 154 | −5,731,800 | 283 | −3,533 | 437 | −5,593,952 | 178 | −5,590,719 | 319 | −3,233 | 497 | −5,455,349 | 201 | −5,452,235 | 326 | −3,115 | 527 |
| LJ-SR-N-N | 832,551 | 45 | 832,706 | 41 | −155 | 85 | 799,353 | 45 | 799,517 | 33 | −164 | 77 | 767,802 | 47 | 767,936 | 39 | −134 | 85 |

[a] Energy terms issued by GROMACS (from the .edr file) in simulations of apoFld with Charmm22-CMAP[1] force field and Tip3p[6] water model.

[b] Averages (Ave) and standard deviations (SD) calculated from the forty (40) replicas simulated for the protein folded (FOLD) state (reported in Galano-Frutos et. al.)[3].

[c] Averages (Ave) and standard deviations (SD) calculated from the hundred (100) unfolded structures simulated for the protein unfolded (UNF) state (reported in Galano-Frutos et. al.)[3].

**SI Table 6.** Data of energy terms (Charmm22-CMAP force field) extracted from the MD simulations of CI2 performed at three temperatures

| | 320 K | | | | | | 335 K | | | | | | 350 K | | | | | |
|---|---|---|---|---|---|---|---|---|---|---|---|---|---|---|---|---|---|---|
| **Force field terms[a]** | **FOLD[b]** | | **UNF[c]** | | **FOLD-UNF[d]** | | **UNF[c]** | | **FOLD[b]** | | **FOLD-UNF[d]** | | **FOLD[b]** | | **UNF[c]** | | **FOLD-UNF[d]** | |
| | **Ave** | **SD** | **Ave** | **SD** | **Ave** | **SD** | **Ave** | **SD** | **Ave** | **SD** | **Ave** | **SD** | **Ave** | **SD** | **Ave** | **SD** | **Ave** | **SD** |
| U-B | 2,587 | 8 | 2,590 | 22 | −3 | 30 | 2,676 | 9 | 2,683 | 23 | −7 | 32 | 2,764 | 10 | 2,770 | 21 | −6 | 31 |
| Proper-Dih | 1,463 | 10 | 1,494 | 32 | −31 | 42 | 1,483 | 10 | 1,523 | 32 | −40 | 41 | 1,510 | 13 | 1,551 | 25 | −41 | 38 |
| Improper-Dih | 143 | 1 | 143 | 1 | 0 | 1 | 149 | 1 | 150 | 1 | 0 | 2 | 156 | 1 | 157 | 1 | −1 | 2 |
| CMAP-Dih | −295 | 7 | −346 | 37 | 52 | 44 | −294 | 8 | −347 | 36 | 53 | 43 | −297 | 10 | −349 | 33 | 52 | 43 |
| LJ-14 | 753 | 3 | 722 | 9 | 31 | 12 | 757 | 3 | 727 | 9 | 30 | 13 | 761 | 4 | 733 | 8 | 28 | 11 |
| Coul-14 | 9,742 | 56 | 9,894 | 86 | −152 | 142 | 9,742 | 56 | 9,884 | 77 | −142 | 134 | 9,768 | 72 | 9,900 | 70 | −132 | 142 |
| LJ-SR | 179,031 | 13 | 179,177 | 22 | −146 | 35 | 171,900 | 14 | 172,051 | 25 | −151 | 39 | 165,087 | 15 | 165,236 | 23 | −149 | 38 |
| Disp-corr | −5,377 | 0 | −5,377 | 0 | 0 | 1 | −5,293 | 0 | −5,293 | 0 | 0 | 0 | −5,204 | 0 | −5,204 | 0 | 0 | 1 |
| Coul-SR | −1,274,111 | 34 | −1,274,174 | 49 | 64 | 83 | −1,242,621 | 37 | −1,242,666 | 47 | 46 | 84 | −1,211,591 | 44 | −1,211,610 | 43 | 20 | 87 |
| Coul-recip | −129,076 | 23 | −129,102 | 38 | 26 | 61 | −128,483 | 24 | −128,504 | 33 | 22 | 57 | −127,867 | 30 | −127,881 | 33 | 14 | 63 |
| Potential | −1,215,140 | 25 | −1,214,979 | 32 | −161 | 57 | −1,189,983 | 30 | −1,189,792 | 31 | −190 | 61 | −1,164,912 | 31 | −1,164,698 | 31 | −213 | 62 |
| Kin-Energy | 249,891 | 3 | 249,891 | 4 | 0 | 7 | 261,605 | 4 | 261,606 | 4 | 0 | 7 | 273,319 | 4 | 273,320 | 4 | −1 | 8 |
| Total-Energy | −965,249 | 24 | −965,088 | 31 | −161 | 56 | −928,378 | 29 | −928,187 | 30 | −191 | 59 | −891,593 | 31 | −891,378 | 31 | −215 | 61 |
| Temp | 320 | 0 | 320 | 0 | 0 | 0 | 335 | 0 | 335 | 0 | 0 | 0 | 350 | 0 | 350 | 0 | 0 | 0 |
| Press | 1 | 0 | 1 | 0 | 0 | 1 | 1 | 0 | 1 | 0 | 0 | 0 | 1 | 0 | 1 | 0 | 0 | 1 |
| Vol | 968 | 0 | 968 | 0 | 0 | 0 | 983 | 0 | 983 | 0 | 0 | 0 | 1,000 | 0 | 1,000 | 0 | 0 | 0 |
| Density | 970 | 0 | 970 | 0 | 0 | 0 | 955 | 0 | 955 | 0 | 0 | 0 | 939 | 0 | 939 | 0 | 0 | 0 |
| pV | 58 | 0 | 58 | 0 | 0 | 0 | 59 | 0 | 59 | 0 | 0 | 0 | 60 | 0 | 60 | 0 | 0 | 0 |
| Enthalpy | −965,191 | 24 | −965,030 | 31 | −161 | 56 | −928,319 | 29 | −928,128 | 30 | −191 | 59 | −891,533 | 31 | −891,318 | 31 | −215 | 62 |
| Coul-SR-P-P | −3,236 | 72 | −2,421 | 157 | −815 | 230 | −3,244 | 85 | −2,472 | 178 | −771 | 264 | −3,285 | 90 | −2,534 | 148 | −751 | 238 |
| LJ-SR-P-P | −1,538 | 13 | −976 | 76 | −562 | 88 | −1,527 | 19 | −985 | 76 | −542 | 96 | −1,512 | 19 | −983 | 69 | −528 | 88 |
| Coul-14-P-P | 9,742 | 56 | 9,894 | 86 | −152 | 142 | 9742 | 56 | 9,884 | 77 | −142 | 134 | 9,768 | 72 | 9,900 | 70 | −132 | 142 |

**SI Table 6.** Continuation...

| Force field terms[a] | 295 K | | | | | | 315 K | | | | | | 335 K | | | | | |
|---|---|---|---|---|---|---|---|---|---|---|---|---|---|---|---|---|---|---|
| | FOLD[b] | | UNF[c] | | FOLD-UNF[d] | | UNF[c] | | FOLD[b] | | FOLD-UNF[d] | | FOLD[b] | | UNF[c] | | FOLD-UNF[d] | |
| | Ave | SD | Ave | SD | Ave | SD | Ave | SD | Ave | SD | Ave | SD | Ave | SD | Ave | SD | Ave | SD |
| LJ-14-P-P | 753 | 3 | 722 | 9 | 31 | 12 | 757 | 3 | 727 | 9 | 30 | 13 | 761 | 4 | 733 | 8 | 28 | 11 |
| Coul-SR-P-N | −5,051 | 124 | −6,978 | 294 | 1,927 | 418 | −4,927 | 149 | −6,738 | 355 | 1,811 | 505 | −4,770 | 152 | −6,508 | 283 | 1,738 | 435 |
| LJ-SR-P-N | −504 | 16 | −957 | 75 | 453 | 91 | −493 | 21 | −918 | 69 | 426 | 91 | −483 | 21 | −890 | 66 | 407 | 87 |
| Coul-SR-N-N | −1,265,825 | 59 | −1,264,776 | 138 | −1,049 | 196 | −1,234,450 | 63 | −1,233,455 | 166 | −994 | 229 | −1,203,536 | 67 | −1,202,569 | 130 | −967 | 197 |
| LJ-SR-N-N | 181,072 | 10 | 181,110 | 16 | −38 | 26 | 173,919 | 9 | 173,954 | 16 | −35 | 25 | 167,082 | 11 | 167,110 | 15 | −28 | 26 |

[a] Energy terms issued by GROMACS (from the .edr file) in simulations of CI2 wild-type with Charmm22-CMAP[1] force field and Tip3p[6] water model.

[b] Averages (Ave) and standard deviations (SD) calculated from the forty (40) replicas simulated for the protein folded (FOLD) state (reported in Galano-Frutos et. al.)[3].

[c] Averages (Ave) and standard deviations (SD) calculated from the hundred (100) unfolded structures simulated for the protein unfolded (UNF) state (reported in Galano-Frutos et. al.)[3].

**SI Table 7.** Data of all energy terms (Amber99SB-ILDN force field) extracted from the MD simulations of barnase performed at three temperatures

| | 305 K | | | | | | 320 K | | | | | | 335 K | | | | | |
|---|---|---|---|---|---|---|---|---|---|---|---|---|---|---|---|---|---|---|
| **Force field terms[a]** | **FOLD[b]** | | **UNF[c]** | | **FOLD-UNF[d]** | | **UNF[c]** | | **FOLD[b]** | | **FOLD-UNF[d]** | | **FOLD[b]** | | **UNF[c]** | | **FOLD-UNF[d]** | |
| | **Ave** | **SD** | **Ave** | **SD** | **Ave** | **SD** | **Ave** | **SD** | **Ave** | **SD** | **Ave** | **SD** | **Ave** | **SD** | **Ave** | **SD** | **Ave** | **SD** |
| Angle | 3,505 | 5 | 3,486 | 26 | 20 | 30 | 3,677 | 9 | 3,670 | 19 | 8 | 28 | 3,852 | 8 | 3,858 | 24 | −6 | 31 |
| Proper-Dih | 4,101 | 6 | 4,069 | 30 | 32 | 36 | 4,163 | 9 | 4,126 | 39 | 37 | 48 | 4,216 | 19 | 4,180 | 29 | 36 | 48 |
| Improper-Dih | 207 | 1 | 205 | 1 | 2 | 2 | 219 | 0 | 217 | 1 | 2 | 2 | 232 | 1 | 230 | 1 | 1 | 2 |
| LJ-14 | 1,861 | 1 | 1,845 | 14 | 16 | 15 | 1,876 | 3 | 1,861 | 12 | 15 | 15 | 1,892 | 4 | 1,878 | 11 | 14 | 16 |
| Coul-14 | 17,964 | 20 | 18,118 | 116 | −154 | 136 | 18,039 | 63 | 18,111 | 130 | −73 | 193 | 18,049 | 68 | 18,111 | 115 | −61 | 183 |
| LJ-SR | 395,771 | 22 | 396,156 | 47 | −385 | 70 | 374,563 | 27 | 374,925 | 58 | −361 | 85 | 354,768 | 23 | 355,129 | 56 | −361 | 79 |
| Disp-corr | −11,295 | 0 | −11,295 | 0 | 0 | 1 | −11,089 | 0 | −11,089 | 0 | 0 | 1 | −10,863 | 0 | −10,862 | 0 | −1 | 1 |
| Coul-SR | −2,739,193 | 38 | −2,739,337 | 108 | 144 | 145 | −2,649,991 | 37 | −2,650,013 | 96 | 22 | 133 | −2,563,127 | 81 | −2,563,051 | 87 | −77 | 168 |
| Coul-recip | −265,724 | 13 | −265,753 | 63 | 29 | 75 | −264,261 | 33 | −264,245 | 74 | −15 | 107 | −262,656 | 38 | −262,633 | 64 | −22 | 101 |
| Potential | −2,592,804 | 24 | −2,592,507 | 47 | −298 | 71 | −2522803 | 31 | −2522436 | 45 | −367 | 76 | −2,453,639 | 50 | −2,453,160 | 45 | −480 | 96 |
| Kin-Energy | 474,543 | 9 | 474,541 | 12 | 2 | 21 | 506,714 | 16 | 506,712 | 14 | 2 | 30 | 538,881 | 21 | 538,886 | 20 | −5 | 41 |
| Total-Energy | −2,118,261 | 19 | −2,117,965 | 53 | −296 | 71 | −2,016,089 | 38 | −2,015,724 | 48 | −365 | 87 | −1,914,757 | 67 | −1,914,273 | 52 | −484 | 119 |
| Temp | 295 | 0 | 295 | 0 | 0 | 0 | 315 | 0 | 315 | 0 | 0 | 0 | 335 | 0 | 335 | 0 | 0 | 0 |
| Press | 1 | 0 | 1 | 0 | 0 | 0 | 1 | 0 | 1 | 0 | 0 | 1 | 1 | 0 | 1 | 0 | 0 | 1 |
| Vol | 1,948 | 0 | 1,948 | 0 | 0 | 0 | 1,984 | 0 | 1,984 | 0 | 0 | 0 | 2,025 | 0 | 2,025 | 0 | 0 | 0 |
| Density | 992 | 0 | 992 | 0 | 0 | 0 | 974 | 0 | 974 | 0 | 0 | 0 | 954 | 0 | 954 | 0 | 0 | 0 |
| pV | 117 | 0 | 117 | 0 | 0 | 0 | 119 | 0 | 119 | 0 | 0 | 0 | 122 | 0 | 122 | 0 | 0 | 0 |
| Enthalpy | −2,118,142 | 20 | −2,117,848 | 52 | −294 | 72 | −2,015,971 | 37 | −2,015,605 | 48 | −366 | 85 | −1,914,635 | 66 | −1,914,152 | 52 | −483 | 119 |
| Coul-SR-P-P | −6,373 | 40 | −4,702 | 209 | −1,671 | 250 | −6,359 | 37 | −4,821 | 229 | −1,537 | 266 | −6,335 | 46 | −4,913 | 202 | −1,423 | 248 |
| LJ-SR-P-P | −3,294 | 14 | −2,046 | 137 | −1,247 | 150 | −3,264 | 15 | −2,097 | 151 | −1,167 | 166 | −3,230 | 21 | −2,120 | 154 | −1,111 | 175 |
| Coul-14-P-P | 17,964 | 20 | 18,118 | 116 | −154 | 136 | 18,039 | 63 | 18,111 | 130 | −73 | 193 | 18,049 | 68 | 18,111 | 115 | −61 | 183 |
| LJ-14-P-P | 1,861 | 1 | 1,845 | 14 | 16 | 15 | 1,876 | 3 | 1,861 | 12 | 15 | 15 | 1,892 | 4 | 1,878 | 11 | 14 | 16 |

**SI Table 7.** Continuation...

| Force field terms[a] | 295 K | | | | | | 315 K | | | | | | 335 K | | | | | |
|---|---|---|---|---|---|---|---|---|---|---|---|---|---|---|---|---|---|---|
| | FOLD[b] | | UNF[c] | | FOLD-UNF[d] | | UNF[c] | | FOLD[b] | | FOLD-UNF[d] | | FOLD[b] | | UNF[c] | | FOLD-UNF[d] | |
| | Ave | SD | Ave | SD | Ave | SD | Ave | SD | Ave | SD | Ave | SD | Ave | SD | Ave | SD | Ave | SD |
| Coul-SR-P-N | −8,419 | 75 | −12,172 | 492 | 3,753 | 567 | −8,296 | 121 | −11,591 | 480 | 3,295 | 602 | −8,146 | 110 | −11,100 | 433 | 2,954 | 544 |
| LJ-SR-P-N | −935 | 10 | −1,895 | 115 | 960 | 125 | −913 | 9 | −1,800 | 123 | 886 | 132 | −889 | 10 | −1,708 | 130 | 819 | 140 |
| Coul-SR-N-N | −2,724,402 | 38 | −2,722,464 | 241 | −1,938 | 279 | −2,635,337 | 94 | −2,633,599 | 232 | −1,738 | 326 | −2,548,646 | 58 | −2,547,038 | 211 | −1,608 | 269 |
| LJ-SR-N-N | 400,000 | 10 | 400,097 | 32 | −98 | 42 | 378,741 | 19 | 378,821 | 30 | −81 | 49 | 358,887 | 14 | 358,956 | 31 | −70 | 45 |

[a] Energy terms issued by GROMACS (from the .edr file) in simulations of barnase with Amber99SB-ILDN[4] force field and Tip3p[6] water model.

[b] Averages (Ave) and standard deviations (SD) calculated from the ten (10) replicas simulated for the protein folded (FOLD) state (reported in Galano-Frutos et. al.[5]).

[c] Averages (Ave) and standard deviations (SD) calculated from the forty (40) unfolded structures simulated for the protein unfolded (UNF) state (reported in Galano-Frutos et. al.[5]).

**SI Table 8.** Data of protein features and experimental values of ΔH and ΔCp of folding

| Protein | PDB ID | Length (no. AAs) | Basic, acidic residues [a] | pI[b] | ΔSASA[c] (Å$^2$) | $\Delta SASA_{pol}$[c] (Å$^2$) | $\Delta SASA_{apol}$[c] (Å$^2$) | Exp $\Delta H_{fold}$[d] (kJ/mol) | Exp $\Delta Cp_{fold}$[e] (kJ/mol·K) |
|---|---|---|---|---|---|---|---|---|---|
| barnase | 1A2P | 110 | 16, 12 | 8.5 | −6,655.1 | −2,182.7 | −4,472.4 | −449.8 | −5.9 |
| SNase | 2SNS | 149 | 32, 19 | 8.9 | −9,167.0 | −3,408.0 | −5,759.0 | −248.2 | −9.6 |
| apoFld | 1FTG | 168 | 15, 31 | 4.0 | −11,474.9 | −3,360.6 | −8,114.3 | −330.5 | −12.6 |
| CI2 | 2CI2 | 65 | 10, 11 | 5.3 | −3,663.3 | −1,261.9 | −2,401.4 | −204.9 | −2.9 |

[a] Basic residues accounted from Lys, Arg and His, and acidic residues accounted from Asp and Glu.

[b] Isoelectric point calculated by the 'IPC protein' method implemented in the Isoelectric Point Calculator server (http://isoelectric.org/).[7]

[c] Solvent-accessible surface area (SASA) change upon folding calculated by the ProtSA server[8] using the referred PDB file and 2000 unfolded structures.[8]

[d] Experimental enthalpy change upon folding extrapolated at the temperatures used for comparison in the main manuscript. Experimental data reported as averages in **Table 1** of Galano-Frutos et. al.[3] used for extrapolating from $\Delta Cp_{fold} = f(\Delta H_{fold}(T))$ linear relationship.

[e] Experimental heat capacity change upon folding as reported in **Table 1** of Galano-Frutos et. al[3].

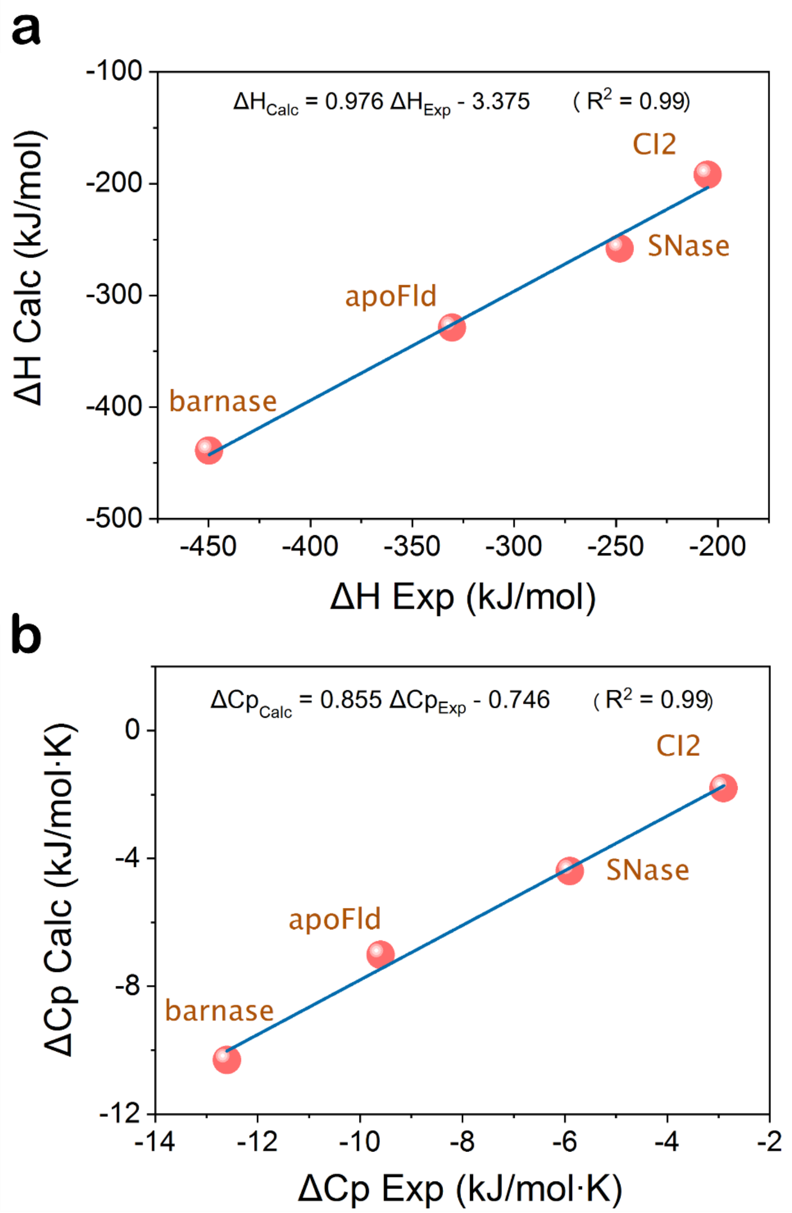


**SI Figure 1. Linear correlation between calculated and experimentally determined $\Delta H_{fol}$ (a) and $\Delta Cp_{fol}$ (b).** Fitting equation and squared Pearson correlation coefficient are included.

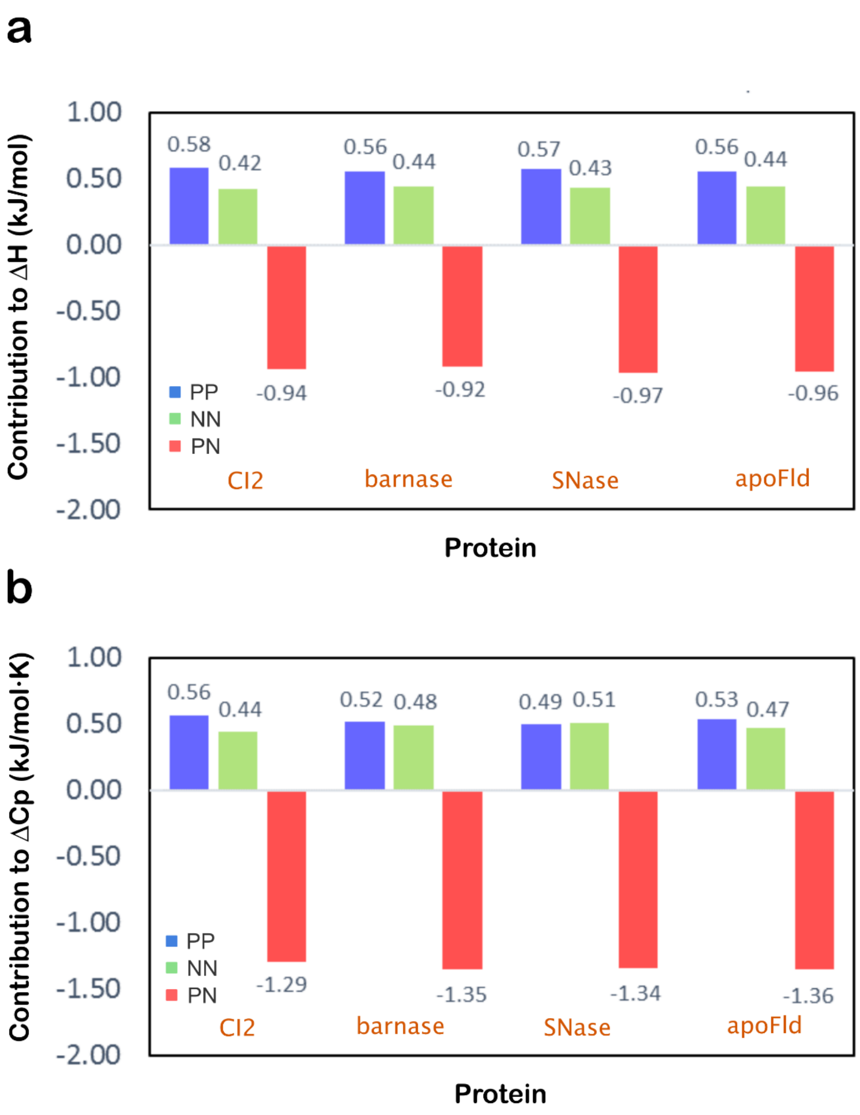


**SI Figure 2. Relative contributions of molecular interactions to $\Delta H_{fol}$ (a) and $\Delta Cp_{fol}$ (b).** 'PP', 'NN' and 'PN' refer to intraproteic, intrasolvent and protein-solvent energy contributions, respectively. Positive contributions add up to 1.

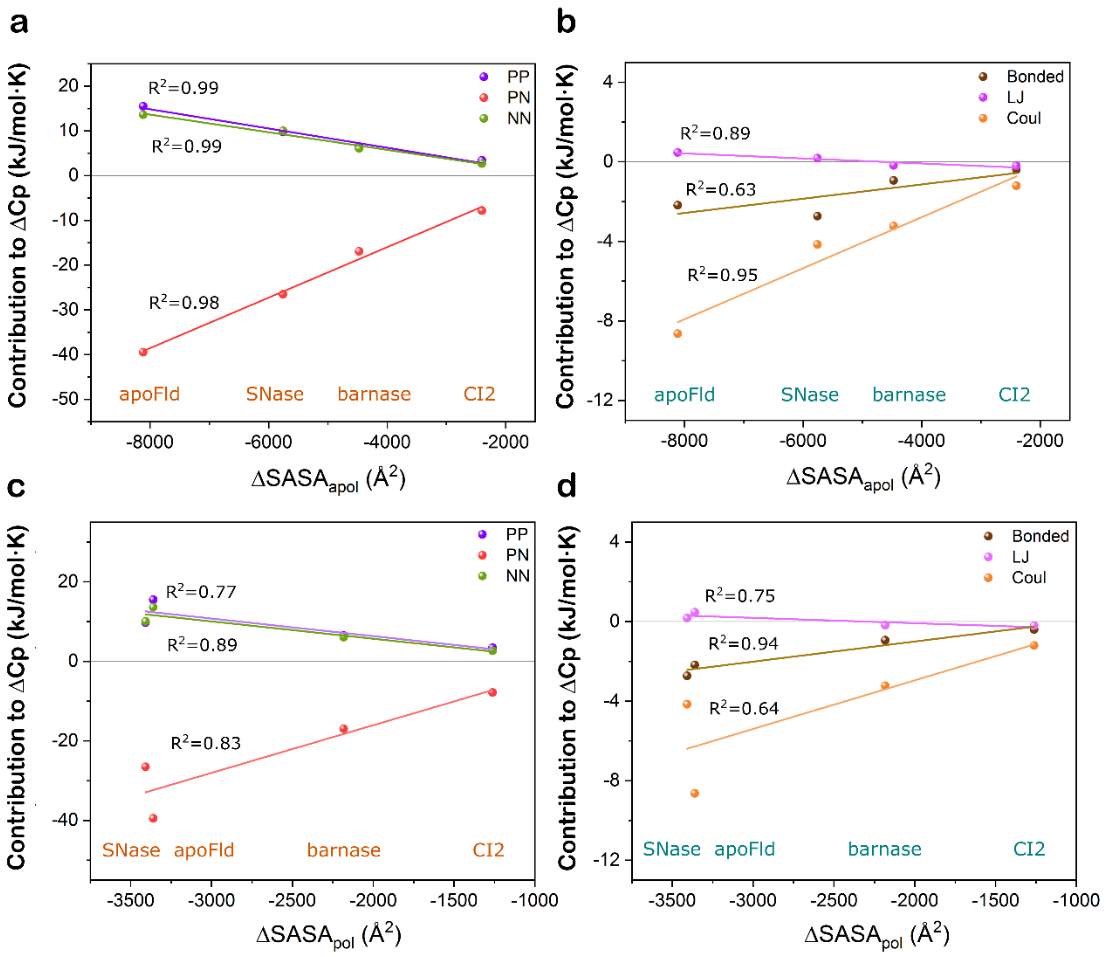


**SI Figure 3. Correlation between $\Delta SASA_{apol}$ (a, b) and $\Delta SASA_{pol}$ (c, d) and molecular (a, c) and elementary (b, d) contributions to $\Delta Cp_{fol}$.** ‘PP’, ‘NN’ and ‘PN’ in **a** and **c** refer to intraproteic, intrasolvent and protein-solvent energy contributions, respectively, whereas ‘Bonded’, ‘LJ’ and ‘Coul’ in **b** and **d** refers to bonded, Lennard-Jones (van der Waals) and coulombic (electrostatics) contributions, respectively. Squared Pearson correlation coefficients are included close to each fitting line.

#### SI References


1. Mackerell, A. D., Feig, M. & Brooks, C. L. Extending the treatment of backbone energetics in protein force fields: Limitations of gas-phase quantum mechanics in reproducing protein conformational distributions in molecular dynamics simulation. *J. Comput. Chem.* **25**, 1400–1415 (2004).
2. Van Der Spoel, D. *et al.* GROMACS: Fast, flexible, and free. *Journal of Computational Chemistry* vol. 26 1701–1718 (2005).
3. Galano-Frutos, J. J., Sancho, J. & Nerín-Fonz, F. Calculation of Protein Folding Thermodynamics using Molecular Dynamics Simulations. *bioRxiv* Preprint (2023) doi:https://doi.org/10.1101/2023.01.21.525008.
4. Lindorff-Larsen, K. *et al.* Improved side-chain torsion potentials for the Amber ff99SB protein force field. *Proteins Struct. Funct. Bioinforma.* **78**, 1950–1958 (2010).
5. Galano-Frutos, J. J. & Sancho, J. Accurate Calculation of Barnase and SNase Folding Energetics Using Short Molecular Dynamics Simulations and an Atomistic Model of the Unfolded Ensemble: Evaluation of Force Fields and Water Models. *J. Chem. Inf. Model.* **59**, 4350–4360 (2019).
6. Jorgensen, W. L., Chandrasekhar, J., Madura, J. D., Impey, R. W. & Klein, M. L. Comparison of simple potential functions for simulating liquid water. *J. Chem. Phys.* **79**, 926–935 (1983).
7. Kozlowski, L. P. IPC - Isoelectric Point Calculator. *Biol. Direct* **11**, 1–16 (2016).
8. Estrada, J., Bernadó, P., Blackledge, M. & Sancho, J. ProtSA: A web application for calculating sequence specific protein solvent accessibilities in the unfolded ensemble. *BMC Bioinformatics* **10**, 1–8 (2009).
